# Subsynaptic positioning of AMPARs by LRRTM2 controls synaptic strength

**DOI:** 10.1101/2021.04.30.441835

**Authors:** A.M. Ramsey, A.H. Tang, T.A. LeGates, X.Z. Gou, B.E. Carbone, S.M. Thompson, T. Biederer, T.A. Blanpied

**Affiliations:** Department of Physiology, University of Maryland School of Medicine, Baltimore, MD, USA; Program in Neuroscience, University of Maryland School of Medicine, Baltimore, MD, USA; Department of Psychiatry, University of Maryland School of Medicine, Baltimore, MD, USA; CAS Key Laboratory of Brain Function and Disease and Hefei National Laboratory for Physical Sciences at the Microscale, School of Life Sciences, Division of Life Sciences and Medicine, University of Science and Technology of China, Hefei, China; Department of Neurology, Yale School of Medicine, New Haven, CT

**Author notes:** Department of Biological Sciences, University of Maryland, Baltimore County, Baltimore, MD, USA. Ohio State University College of Medicine, Columbus, OH, USA. These authors contributed equally (A.M. Ramsey, A.H. Tang).

## Abstract

Recent evidence suggests that nanoorganization of proteins within synapses may control the strength of communication between neurons in the brain. The unique subsynaptic distribution of glutamate receptors, which cluster in nanoalignment with presynaptic sites of glutamate release, supports this idea. However, testing it has been difficult because mechanisms controlling subsynaptic organization remain unknown. Reasoning that transcellular interactions could position AMPA receptors, we targeted a key transsynaptic adhesion molecule implicated in controlling AMPAR number, LRRTM2, using engineered, rapid proteolysis. Severing the LRRTM2 extracellular domain led quickly to nanoscale de-clustering of AMPARs away from release sites, not prompting their escape from synapses until much later. This rapid remodeling of AMPAR position produced significant deficits in evoked, but not spontaneous, postsynaptic receptor activation. These results dissociate receptor numbers from their nanopositioning in determination of synaptic function, and support the novel concept that adhesion molecules acutely position AMPA receptors to dynamically control synaptic strength.

## Introduction

The complex neural processes of information encoding, storage, and retrieval are enabled by fine regulation of synaptic strength. It is well established that the number of AMPA-type glutamate receptors (AMPARs) within a single synapse is a key property determining the amplitude of the excitatory postsynaptic current (EPSC) response to neurotransmitter release (*1–3*). However, several constraints appear to limit AMPAR activation following glutamate release (*4–6*), suggesting that factors beyond receptor number also control synapse strength. Among the most critical is that a variety of modeling approaches suggest AMPARs even ~90 nm from the site of glutamate release open with only about half the likelihood of those close to the site of vesicle fusion (*7*, *8*), due to the low affinity and rapid desensitization of receptors (*9–11*). This greatly diminishes the expected EPSC, a prediction in line with experimental results suggesting that glutamate release from a single vesicle is not sufficient to maximize postsynaptic receptor activation (*12*, *13*). Unfortunately, it has been difficult to test whether such distance-dependence plays a physiological role in neurons because the mechanisms that determine the precise positioning of receptors across from sites of release are not known.

Discerning these mechanisms is complex because AMPARs and a number of scaffolding molecules involved in their synaptic retention, most notably PSD-95, are non-homogenously distributed within individual PSDs, forming nanometer-scale clusters (*14–17*). Similarly, within the presynaptic active zone (AZ), molecules critical for vesicle priming and Ca2+ channel recruitment such as the Rab3 Interacting Molecule (RIM) and Munc13 are clustered into ~100 nm nanodomains (*18*). These AZ nanodomains are widely conserved across many synapse types (*19*) and are thought to govern vesicle positioning and establish sites in the AZ where action potentials drive synaptic vesicle exocytosis with highest probability (*20*, *21*). Critically, in mammalian brain, presynaptic sites of glutamate exocytosis as marked by RIM nanoclusters are aligned with postsynaptic nanoclusters of AMPARs across the cleft in an organization referred to as a nanocolumn (*20*, *22*, *23*). If receptor distance to the site of neurotransmitter exocytosis regulates receptor activation, then this aligned organization likely enhances basal excitatory synaptic transmission, and its disruption would reduce synaptic strength. This is important to determine, since modulation of transsynaptic alignment then would open a number of different mechanisms of synaptic plasticity (*24*).

It remains unclear how subsynaptic alignment of receptor clusters with release sites is created or maintained. Though many models have been proposed (*5*), perhaps the most parsimonious idea is that cleft-resident synaptic cell adhesion molecules (CAMs) link pre- and postsynaptic nanodomains through their transsynaptic binding interactions. The most prominent candidates to test for this role are the neuroligin (NL) and Leucine Rich Repeat Transmembrane (LRRTM) families that are postsynaptic partners of presynaptic neurexins and that bind PSD-95 (*25*). However, such tests are complicated because CAM families are large, the roles they play are diverse, and the family members exhibit substantial redundancy upon knockout (*26*, *27*). Indeed, disruption of postsynaptic NL by expression of dominant negative mutants or prolonged incubation with interfering peptides does in fact alter receptor alignment with RIM (*28*), providing support for the idea. However, these extended treatments also prompt a complex set of other effects including altering synapse numbers, presynaptic vesicle release probability, and frequency of spontaneous transmission (*28–31*).

The LRRTM family are strong candidates to mediate transsynaptic alignment. A key abundant family member in hippocampus, LRRTM2, binds postsynaptic PSD-95 through a C-terminal motif (*32*, *33*) and the presynaptic Neurexin-Heparan sulfate complex through 10 extracellular LRR repeats (*34*). LRRTM2 has been found to be important for evoked AMPAR-mediated, though not NMDAR-mediated, synaptic transmission independent of synaptogenesis (*35*). Furthermore, the extracellular domain of LRRTM2 alone is sufficient to rescue AMPAR-mediated synaptic transmission following LRRTM1,2 double knockout, a mechanism proposed to be achieved by the anchoring of AMPARs in the PSD (*35*, *36*). LRRTM2 within synapses also forms nanoscale clusters of similar size to scaffold, receptor, and release machinery nanodomains (*37*). Thus, we hypothesized that LRRTM2 coordinates positioning of receptors relative to evoked release sites.

Long-term manipulations can prompt substantial reorganization of synapses which makes deducing the native state difficult. To test the role of LRRTM2 while avoiding such effects, we used acute extracellular proteolysis of an engineered cleavage site to disrupt its extracellular interactions within seconds, thus uncoupling it from the postsynaptic membrane while avoiding complications of genetic compensation. With this approach, we discovered that LRRTM2 acutely controls the fine positioning of AMPARs relative to the site of release. The repositioning of AMPARs following loss of the LRRTM2 extracellular domain leads to reduction in the amplitude of evoked but not spontaneous responses. Further, the basal distribution of LRRTM2 is in nanoscale register with both RIM and AMPAR nanodomains. Together, these data suggest that postsynaptic LRRTM2 establishes a transcellular, structural linkage mediating nanocolumn alignment of AMPARs with preferential sites of evoked neurotransmitter release and provide strong evidence that AMPAR organization within the synapse is critical for the strength of basal synaptic transmission.

## Results

### Acute and specific cleavage of the LRRTM2 extracellular domain

To test the role of LRRTM2 extracellular interactions in synapse structure and function independent of synaptogenesis and genetic compensation, we adapted a previous approach (*38*) and inserted the short recognition sequence for the endoprotease thrombin (LVPRGS) at an extracellular, juxtamembrane position within human LRRTM2 (Fig. 1A). To visualize the molecule and enable live-cell measurement of its cleavage in neurons, we appended EGFP to the N-terminus, and used this to replace endogenous LRRTM2 following knockdown with published shRNA targeting sequences. We denote the molecule GFP-Thr-LRRTM2*, where * indicates the human sequence designed to be resistant to the shRNA (*33*, *39*).

**Fig. 1.**
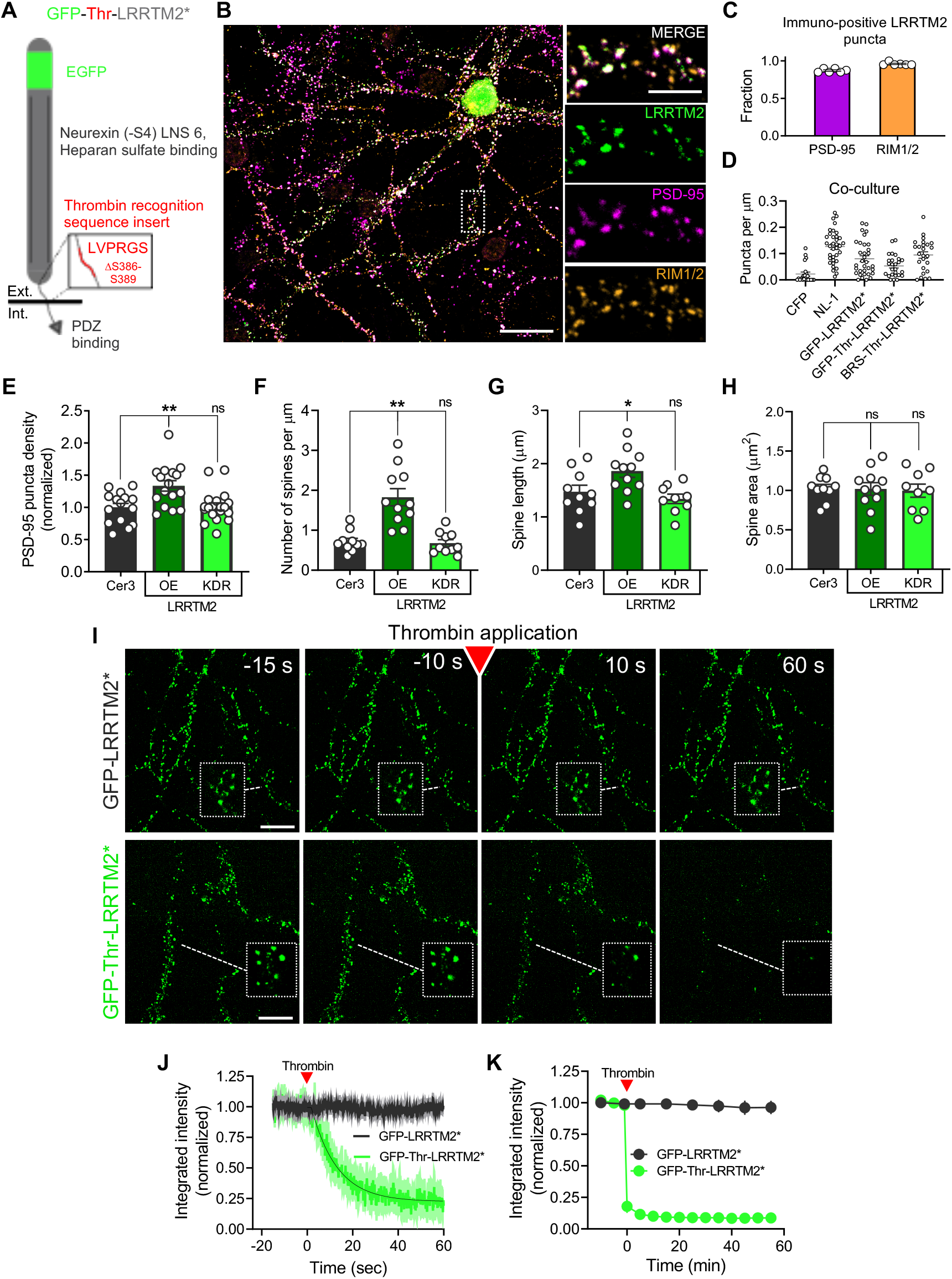
Acute and specific cleavage of the LRRTM2 extracellular domain. **(A)** Schematic demonstrating the juxtamembrane insertion of the thrombin recognition sequence (*38*) and the N-terminal GFP* denotes co-packaging of an shRNA (*33*) that targets endogenous LRRTM2 expressed in the same vector as GFP-Thr-LRRTM2. **(B)** Expression of GFP-Thr-LRRTM2* in cultured hippocampal neurons and immunostaining of endogenous PSD-95 and RIM1/2 visualized by confocal microscopy. Scale bar, left: 30 μm, right: 10 μm. **(C)** Quantification of the colocalization between expressed GFP-Thr-LRRTM2*, RIM1/2, and PSD-95. (*n* = 120 synapses/6 neurons/2 independent cultures per condition). **(D)** Quantification of Bassoon recruitment by LRRTM2 in an HEK-neuron coculture synaptogenesis assay alongside positive (CFP-NL1) and negative (CFP alone) controls. CFP alone (*n* = 30 cells/2 independent cultures), CFP-NL1 (*n* = 34/2), BRS-Thr-LRRTM2* (*n* = 24/2), GFP-Thr-LRRTM2* (*n* = 25/2), GFP-LRRTM2* (*n* = 32/2). **(E)** Quantification of PSD-95 puncta density in neurons expressing GFP-Thr-LRRTM2* (KDR, *n* = 19 neurons/3 independent cultures), GFP-Thr-LRRTM2 (OE, *n* = 16/3), or cytosolic mCerulean3 (Cer3, *n* = 15 /3). **(F)** Quantification of spine density. (*n* = 10 neurons/3 independent cultures per condition). **(G)** Quantification of spine length. (*n* = 10/3). **(H)** Quantification of spine area. (*n* = 10/3). **(I)** Representative images from a confocal time series of GFP-Thr-LRRTM2* cleavage following thrombin application (red arrow, 10 units ml^-1^). Scale bar: 10 μm. **(J)** Quantification of GFP-Thr-LRRTM2* (*n* = 100 synapses/5 neurons/3 independent cultures) and GFP-LRRTM2* (*n* = 40/2/2) cleavage. **(K)** Quantification of GFP-LRRTM2* (*n* = 100/5/2) or GFP-Thr-LRRTM2* (*n* = 120/6/2) for up to 60 minutes post thrombin exposure. One-way ANOVA with posthoc Dunnett’s Test was used in E-H. Data are presented as mean ± SEM, * *p* ≤ 0.05, ** *p* ≤ 0.01.

These modifications of LRRTM2 did not appear to disrupt its function. When expressed in cultured rat hippocampal neurons, GFP-Thr-LRRTM2* clustered avidly in small puncta that colocalized nearly exclusively with synapses immunolabeled for PSD-95 and RIM1/2 (Fig. 1B-C), though some puncta appeared in the dendritic shaft apart from synapses. In addition, when expressed in HEK cells, GFP-Thr-LRRTM2* trafficked strongly to the plasma membrane and retained the synaptogenic ability of wild type LRRTM2 to cluster presynaptic markers in the axons of co-cultured wild type neurons (Fig 1D, Supplementary Fig. 1A).

Though knockdown of LRRTM2 was successful (Supplementary Fig. 2), typical rescue strategies can still result in overexpression. Since overexpression of LRRTM2 in cultured neurons increases excitatory synapse density (*33*, *39*), we tested for functional effects of LRRTM2 overexpression by measuring synapse density via PSD-95 immunolabeling. As expected, expression of GFP-Thr-LRRTM2* without concurrent knockdown resulted in a ~1.3-fold increase in PSD-95 puncta compared to controls expressing cytosolic mCerulean3 alone (Fig. 1E). However, puncta density was unchanged following knockdown of endogenous LRRTM2 and replacement with GFP-Thr-LRRTM2*. Similarly, compared to mCerulean transfected neurons, spine numbers were increased by GFP-Thr-LRRTM2* overexpression, but were unchanged following knockdown and replacement (Fig. 1F). Overexpression also resulted in an increase in spine length though not spine area, but we found no changes in either measure with the knockdown-replacement approach (Fig. 1G-H). The replacement strategy also minimized non-synaptic localization of GFP-Thr-LRRTM2*, which was enriched much more specifically in synapses when endogenous LRRTM2 was knocked-down as judged by the levels of thrombin-sensitive extrasynaptic fluorescence (Supplementary Fig. 1B). These data suggest that GFP-Thr-LRRTM2* incorporates readily into excitatory synapses without disrupting synaptogenesis and with minimal effects of overexpression.

Next, we tested whether thrombin successfully cleaved GFP-Thr-LRRTM2* in synapses, particularly at low working concentrations to avoid potential effects which could be mediated via PAR receptors (*40*). Following baseline measurements of EGFP fluorescence, we bath-applied thrombin at 10 units/mL. This prompted the rapid and robust loss of GFP fluorescence from puncta in dendritic spines (Fig. 1I, below). Thrombin application to neurons expressing GFP-LRRTM2* (with no thrombin recognition sequence) resulted in no decrease in fluorescence, indicating the loss was due to cleavage of the extracellular domain (ECD) and not non-specific effects of thrombin (Fig. 1I, above). The LRRTM2 ECD was lost with a time constant of τ = 11.08 seconds (95% C.I 10.74 to 11.43 sec; Fig. 1J), surprisingly rapid given its presumed interactions within the synaptic cleft. Incubations in thrombin for up to 1 hour showed sustained loss of GFP-Thr-LRRTM2* (Fig 1K; fractional fluorescence remaining; 0.09 ± 0.02 compared to baseline; mean ± SEM), but no loss of the LRRTM2 ECD in GFP-LRRTM2* transfected neurons (Fig 1K; fraction remaining; 0.96 ± 0.05, compared to baseline). The rate of cleavage is likely limited by the speeds of perfusion and proteolysis, but regardless suggests that LRRTM2 ECD interactions are insufficient to immobilize it for substantial periods within the synaptic cleft. In addition, the quick action and extensive loss of fluorescence confirms that GFP-Thr-LRRTM2* was trafficked to the cell surface as expected, and suggests that LRRTM2 is only minimally retained intracellularly at steady state in these neurons. Overall, these results demonstrate that expressed GFP-Thr-LRRTM2* localizes appropriately to excitatory synapses, retains its synaptogenic activity, induces no observable morphological changes in spines, and can be proteolytically cleaved acutely and specifically on demand.

### No rapid loss of AMPARs following removal of the LRRTM2 extracellular domain

The role of LRRTM2 in synaptogenesis has been well studied, but its functions in established synapses have been explored in far less detail. The four C-terminal amino acids in its intracellular domain form a PDZ-binding motif which is thought to play a role in the recruitment of PSD-95 in developing synapses (*33*), through interactions with the first and second PDZ domains of PSD-95 (*32*). We considered whether maintenance of PSD-95 at established synapses depends on stable LRRTM2 extracellular interactions. To test this, we co-transfected neurons with GFP-Thr-LRRTM2* and PSD95*-mCherry (*15*) (here, * also denotes resistance to co-expressed shRNA) and measured their fluorescence intensity over the course of a 30 min thrombin application. Strikingly, despite a large and sustained reduction in the GFP-Thr-LRRTM2* fluorescence (fraction remaining: 0.15 ± 0.07, Fig. 2A-C), PSD95*-mCherry fluorescence at synapses remained unchanged (fraction remaining: 0.94 ± 0.05, Fig. 2A-C). Immunocytochemical analysis of synaptic PSD-95 content after thrombin cleavage (discussed below) confirmed this result. Thus, the interactions of LRRTM2 within the synaptic cleft are not necessary for the retention of PSD-95 in established synapses.

**Fig. 2.**
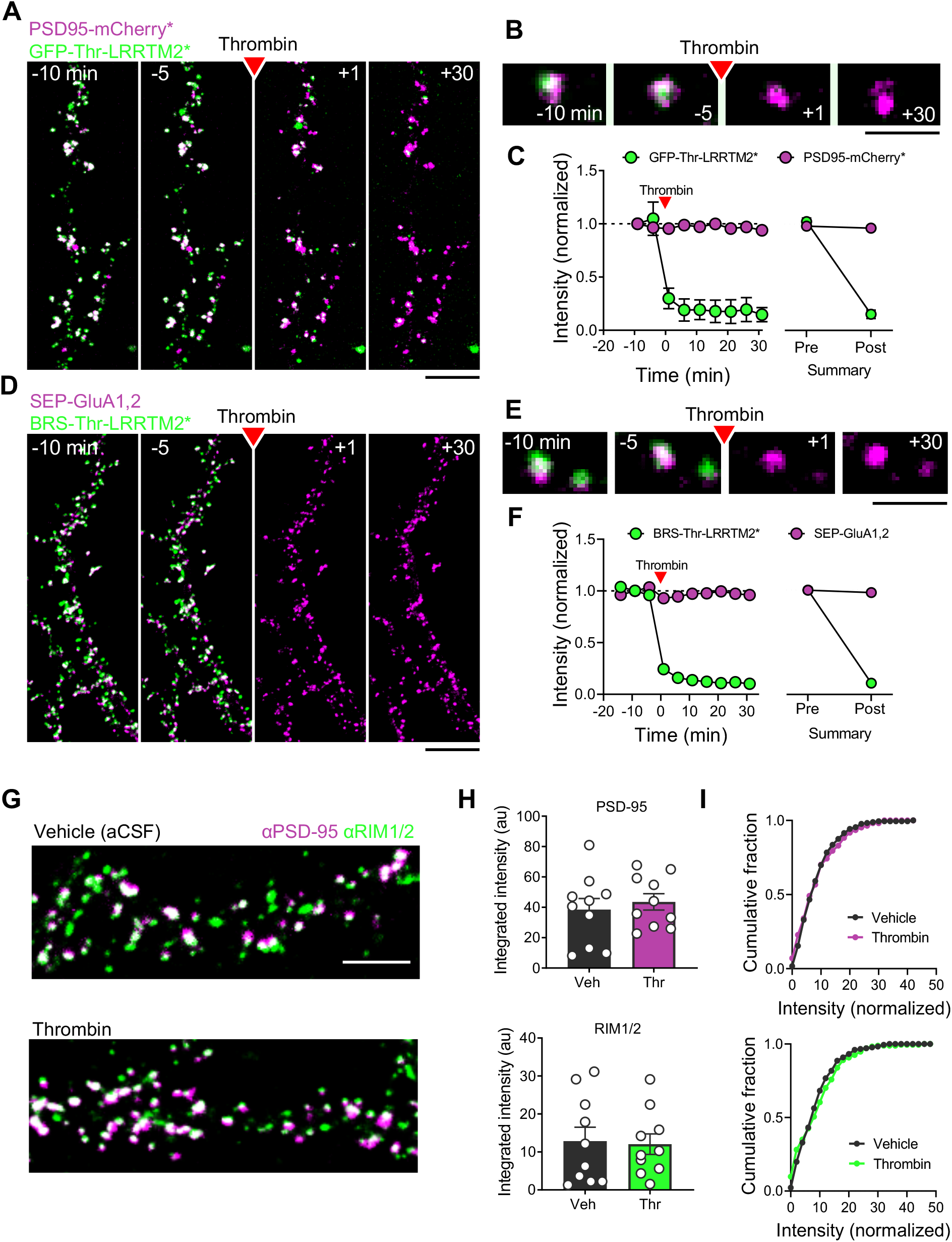
No rapid loss of AMPARs following removal of the LRRTM2 extracellular domain. **(A)** Representative images of neuronal dendrites co-expressing GFP-Thr-LRRTM2* and PSD95-mCherry*. Red arrow indicates the bath application of thrombin (10 units ml^-1^). Scale bar: 10 μm. **(B)** Enlarged view. Scale bar: 2 μm. **(C)** Left, quantification of fluorescence intensity of both GFP-Thr-LRRTM2* and PSD95-mCherry*. Right, summary of baseline measurements compared to 30’ post thrombin application. (*n* = 14 neurons/3 independent cultures). **(D)** Representative images of neuronal dendrites co-expressing BRS-Thr-LRRTM2* and SEP-GluA1/2. Red arrow indicates the bath application of thrombin (10 units ml^-1^). Scale bar: 10 μm. **(E)** Enlarged view. Scale bar: 2 μm. **(F)** Left, quantification of fluorescence intensity of both BRS-Thr-LRRTM2* labeled with α-bungarotoxin conjugated to Alexa-647 and SEP-GluA1/2 over time normalized to their respective baseline. Right, summary of baseline measurements compared to 30’ post thrombin application. (*n* = 11 neurons/3 independent cultures). **(G)** Representative images of immunocytochemical staining of endogenous RIM1/2 and PSD-95 from cultured hippocampal neurons expressing GFP-Thr-LRRTM2* and mCerulean3 treated with either vehicle (aCSF, above; n =173 synapses/9 neurons/3 independent cultures) or thrombin (below, 10 units ml^-1^ for 10 minutes; n = 176/9/3). Scale bar: 5μm. **(H)** Quantification of synaptic staining intensity for PSD-95 (above) and RIM1/2 (below). **(I)** Cumulative distribution of synaptic staining intensities for cells treated with vehicle (aCSF, grey) or thrombin (magenta for PSD-95 and green for RIM1/2). Data are presented as mean ± SEM.

LRRTM2 is important for both establishing the number of GluA1-containing synapses as well as basal synaptic content of GluA1 and GluA2 (*33*, *36*, *41*). In neurons at rest, many AMPARs continuously diffuse within the synapse and exchange between synaptic and extrasynaptic domains on a time scale of seconds to minutes (*42*), but the mechanisms that counteract diffusion and enrich them in the PSD are incompletely understood. Extracellular interactions in the synaptic cleft may be important, and it is conceivable that the ECD interactions of LRRTM2 could assist in the stabilization of both LRRTM2 and additionally GluA1-containing AMPARs in established synapses. To visualize synaptic AMPAR content during live imaging before and after cleavage of LRRTM2, we utilized super-ecliptic pHluorin (SEP)-tagged GluA1 and GluA2, as previously described (*43*). We expressed these receptors along with a version of LRRTM2* in which the GFP was replaced with the smaller α-bungarotoxin recognition sequence (*44*) (BRS-Thr-LRRTM2*) which retained its synaptogenic activity (see Fig 1D, Supplementary Fig. 1) and allowed us to select the wavelength of the labeled α-bungarotoxin. Alexa-647 conjugated to α-bungarotoxin was applied to live cells, resulting in synapse-specific labeling and visualization of the LRRTM2 ECD in co-transfected neurons. We predicted that if the LRRTM2 ECD interacts directly or indirectly with the GluA1 extracellular domain, its acute loss would reduce SEP-GluA1,2 content in synapses. As with GFP-Thr-LRRTM2*, thrombin application produced a rapid and dramatic loss of α-bungarotoxin-Alexa-647 fluorescence (fraction remaining: 0.10 ± 0.02, Fig. 2D-F) indicating cleavage and dispersal of the LRRTM2 ECD. However, SEP-GluA1,2 fluorescence colocalized with LRRTM2 puncta did not decrease even after 10 or 30 min (fraction remaining: 0.96 ± 0.03, Fig. 2D-F), suggesting no changes in the number of receptors present within the PSD. Furthermore, neurons expressing SEP-GluA1/2 along with the cleavable or non-cleavable versions of LRRTM2*, showed no difference in the SEP-GluA1,2 synaptic cluster localization density as measured by dSTORM after a 10-min thrombin treatment (Mann-Whitney Test, p = 0.85; data discussed below, Supplementary Fig. 3). These data suggest that the LRRTM2 ECD is not required for the synaptic retention of AMPARs within a time frame of 30 minutes, during which many receptors exchange in and out of the synapse (*45*).

This result was surprising because conditional knockout of LRRTM1 and 2 leads to a reduction in AMPAR content and EPSC amplitude at established synapses (*35*). One major difference between the conditional deletion and the acute cleavage is the vastly differing time scales of the two approaches. To test whether the prolonged loss of the LRRTM2 ECD affects AMPAR retention in spines, we performed live-cell imaging for up to 2 hours post-cleavage. In fact, synaptic AMPAR content remained almost completely unchanged for at least 60 min after LRRTM2 cleavage. Only after this, a slow decline set in, and 2 hours after cleavage there was a 23.55% ± 0.08% reduction in AMPAR content compared to non-cleavable controls (Supplementary Fig. 4). To examine longer time points, we fixed transfected neurons 24 hours after thrombin treatment and stained for surface SEP-GluA1,2. Compared to controls expressing non-cleavable BRS-LRRTM2*, neurons that underwent LRRTM2 cleavage displayed much weaker surface SEP-GluA1,2 expression (Supp. Fig. 4). These data corroborate the previously reported idea that LRRTM2 is important for AMPAR stability in synapses (*35*), but show that this effect plays out only over extended periods without the LRRTM2 ECD.

Another possible role of LRRTM2 may be to instruct organization of presynaptic release machinery. To test this, we transfected cultured hippocampal neurons with GFP-Thr-LRRTM2* along with soluble Cerulean3 to identify transfected spines following elimination of the EGFP fluorescence post-cleavage. Then, following live-cell cleavage of LRRTM2 with 10 U/mL thrombin for 10 minutes, cells were fixed, permeabilized, and stained for endogenous PSD-95 and the critical presynaptic scaffolding molecule RIM1/2. Despite near complete elimination of EGFP fluorescence at transfected spines we observed no changes in endogenous RIM1/2 content (fraction remaining: 1.06 ± 0.06, compared to vehicle, Fig. 2G-I). These data suggest that LRRTM2 is not necessary for the retention of RIM in the active zone. Analysis of PSD-95 staining intensity further confirmed that cleavage of LRRTM2 in established synapses did not change PSD-95 content (fraction remaining; 1.02 ± 0.06, compared to vehicle, Fig. 2G-I) supporting our earlier observations during live imaging. Taking these data together, we conclude that although the LRRTM2 ECD is quickly lost after thrombin cleavage in established synapses, its acute removal does not rapidly lead to loss of other key molecules, including AMPARs.

### LRRTM2 is enriched within the trans-synaptic nanocolumn

Growing evidence indicates that different CAMs possess unique and distinct organizations within excitatory synapses (*37*). For instance, both SynCAM1 and Neurexin-1 are enriched in a small number of subsynaptic ensembles, but the nanoclusters of SynCAM-1 are often found near or around the border of the synapse (*46*, *47*) whereas Neurexin-1 nanoclusters tend to occur just slightly off-center within the PSD (*48*). How these distributions subserve particular functions is not known. LRRTM2 forms tight clusters in the postsynaptic density (*37*). Its enrichment within these nanoclusters in notably tighter than Neuroligin-1, which more homogeneously distributes through the synapse (*37*), but neither the location nor function of LRRTM2 nanoclusters is known. We hypothesized that LRRTM2 may link pre- and postsynaptic nanodomains, and therefore predicted that it is enriched with other proteins found within the trans-synaptic nanocolumn (*20*).

To test whether LRRTM2 formed subsynaptic clusters within excitatory PSDs, we performed two-color 3D dSTORM in our LRRTM2 knockdown-replacement system using an anti-GFP antibody (Fig. 3A). Maps of the local density at each molecular location (Fig. 3B) showed that LRRTM2 is non-uniformly organized within the PSD, forming nanodomains of similar size and number to those previously reported for receptors and scaffolding molecules (*14*, *15*, *37*). To quantify the degree to which LRRTM2 was self-clustered, we calculated an autocorrelation measurement and found that LRRTM2 was robustly organized into clusters with a ~100 nm diameter within synapses (Fig. 3C).

**Fig. 3.**
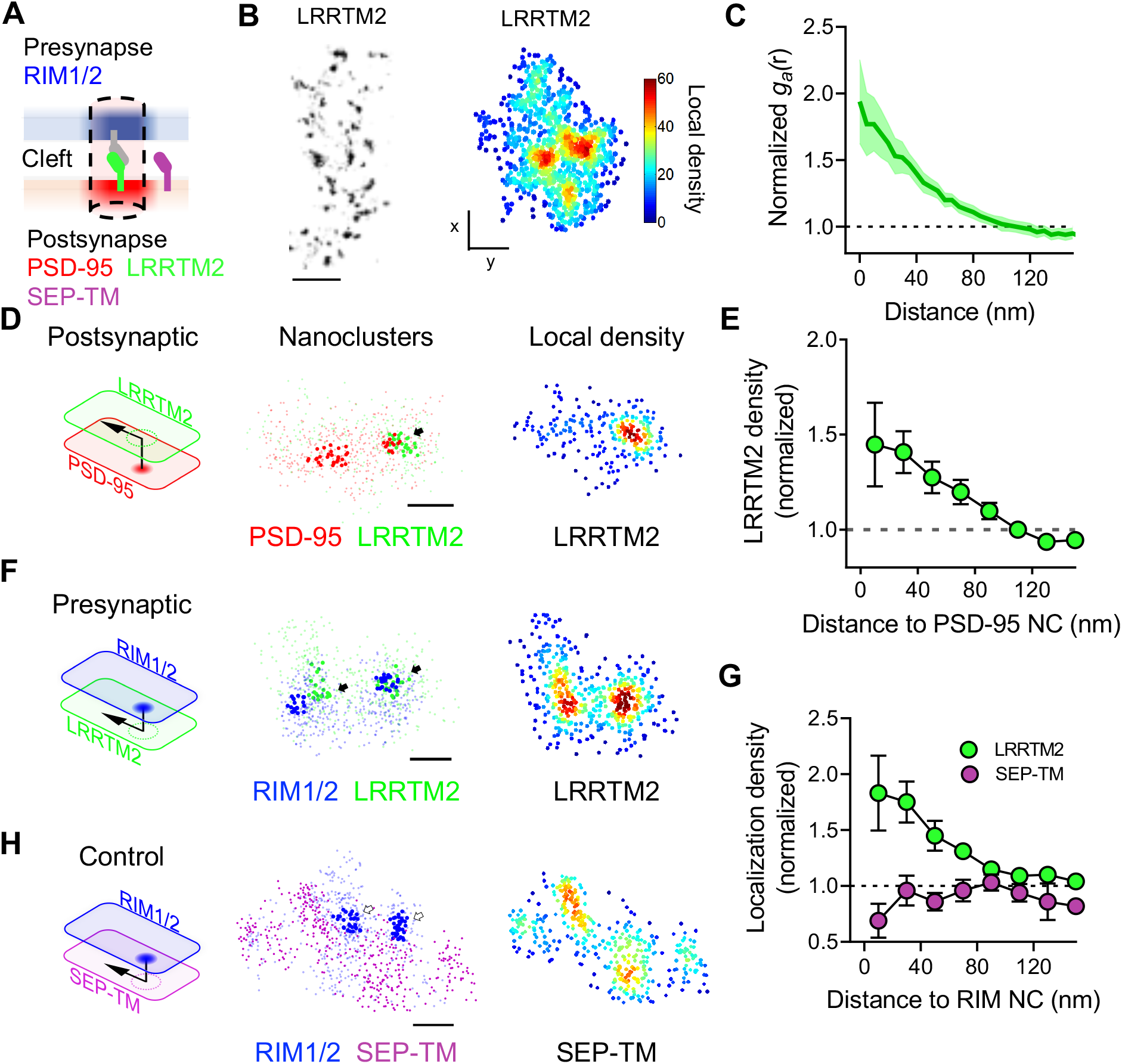
LRRTM2 is enriched within the trans-synaptic nanocolumn. **(A)** Schematic demonstrating the trans-synaptic nanoscale organization of LRRTM2 relative to RIM and PSD-95. **(B)** Left, 3D dSTORM reconstruction of a dendrite from a neuron expressing GFP-LRRTM2*. Scale bar: 2 μm. Right, 3D dSTORM reconstructions of an individual synapse with localizations color-coded by local density (5x nearest neighbor distance (NND)). Scale bar: 100 nm. **(C)** Auto-correlation of LRRTM2 synaptic clusters. **(D)** Schematic demonstrating the measurement of 3D co-enrichment between protein pairs (LRRTM2 green, PSD-95 red). Middle, left, en face view of the localized positions of PSD-95 (red) and LRRTM2 (green) with detected nanoclusters indicated in bold. Middle, right, the same LRRTM2 localizations coded by their local density (5x NND). Scale bar: 100 nm. **(E)** Quantification of LRRTM2 density as a function of the distance to the PSD-95 nanocluster center. **(F)** LRRTM2 cross-enrichment with RIM1/2 as displayed in DE. **(G)** Quantification LRRTM2 cross-enrichment with RIM1/2 nanoclusters. (*n* = 176 nanoclusters/16 neurons/5 independent cultures). **(H)** Cross-enrichment of a diffuse target, SEP-TM, across from presynaptic RIM1/2 nanoclusters displayed as in D-E. Scale bar: 100 nm. (*n* = 85/9/3). **(G)** Quantification of SEP-TM density as a function of the distance to the RIM1/2 nanocluster center. Data are presented as mean ± SEM.

To test how LRRTM2 was organized relative to nanocolumn-resident molecules, we measured the subsynaptic organization of GFP-Thr-LRRTM2* relative to endogenous PSD-95 (Fig. 3D-E) and RIM1/2 (Fig. 3F-G) using an enrichment assay reported previously (*20*). LRRTM2 was tightly enriched within PSD-95 nanodomains (enrichment index: 1.37 ± 0.13, Fig. 3E) and aligned with RIM1/2 nanodomains across the synaptic cleft (enrichment index: 1.56 ± 0.21%, Fig. 3G). To analyze the LRRTM2 distribution with respect to that of RIM1/2 or PSD-95 without requiring the identification of nanoclusters of either protein, we measured the cross-correlation between LRRTM2 and RIM1/2 or PSD-95 density distributions (Sup. Fig. 3). This demonstrated that the distribution of LRRTM2 and PSD-95 as well as LRRTM2 and RIM1/2 are highly similar to one another. To illustrate this similarity, we compared the distribution of LRRTM2 to a probe without nanocolumns enrichment. We used an engineered single pass transmembrane protein called SEP-TM containing an N-terminal extracellular super-ecliptic pHluorin (SEP) appended to the transmembrane domain from PDGFR (*49*). SEP-TM traffics avidly to the plasma membrane (Supplementary Fig. 5), but we predicted that it would not be enriched within the nanocolumn because it lacks relevant binding via its N- or C-terminus. While LRRTM2 was tightly aligned across the cleft from RIM nanodomains, SEP-TM was not (enrichment index: 0.83 ± 0.12, Fig. 3G-H, Supplementary Fig. 6). This suggests that the subsynaptic positioning of LRRTM2 is actively determined by protein-protein interactions rather than arising as a general feature of transmembrane proteins in the dense synaptic environment. The distribution of LRRTM2 within the synapse, tightly clustered and colocalized with transsynaptically aligned protein nanodomains, suggests that it acts within the nanocolumn.

### LRRTM2 is critical for AMPAR enrichment with preferential sites of evoked neurotransmitter release

Expression of LRRTM2 with mutations that disrupt its interaction with presynaptic neurexin decreases content of expressed mutant LRRTM2 at synapses and leads to lower synaptic AMPAR content and reduced AMPAR-mediated EPSCs (*35*, *41*). However, it remains unclear whether LRRTM2 exerts ongoing control of synaptic transmission in established synapses. Given the location of LRRTM2 within the nanocolumn, we reasoned that its extracellular interactions may contribute to the nanoscale alignment of AMPARs to RIM nanodomains. To test this, we took advantage of the acute nature of the protease cleavage approach, which avoids the complications of compensation by other CAMs during the prolonged periods required for molecular expression.

Since we wanted to assure that we measured receptors only in cells where we manipulated LRRTM2 rather than nearby untransfected synapses, we first used two-color 3D dSTORM to measure the distribution of SEP-tagged AMPARs co-transfected with LRRTM2. Our previous work demonstrated that endogenous receptors are enriched in ~80 nm nanodomains aligned with surprising precision to presynaptic RIM nanodomains (*20*). As expected, SEP-GluA1/2 AMPARs in neurons co-transfected with BRS-Thr-LRRTM2* but treated only with vehicle formed nanodomains of ~80 nm, as judged by the autocorrelation of their distributions (Supplementary Fig. 7A,B). These were strongly enriched with RIM nanodomains across the cleft, decaying in enrichment over approximately 80 nm (Fig 4A, Supplementary Fig. 7C).

**Fig. 4.**
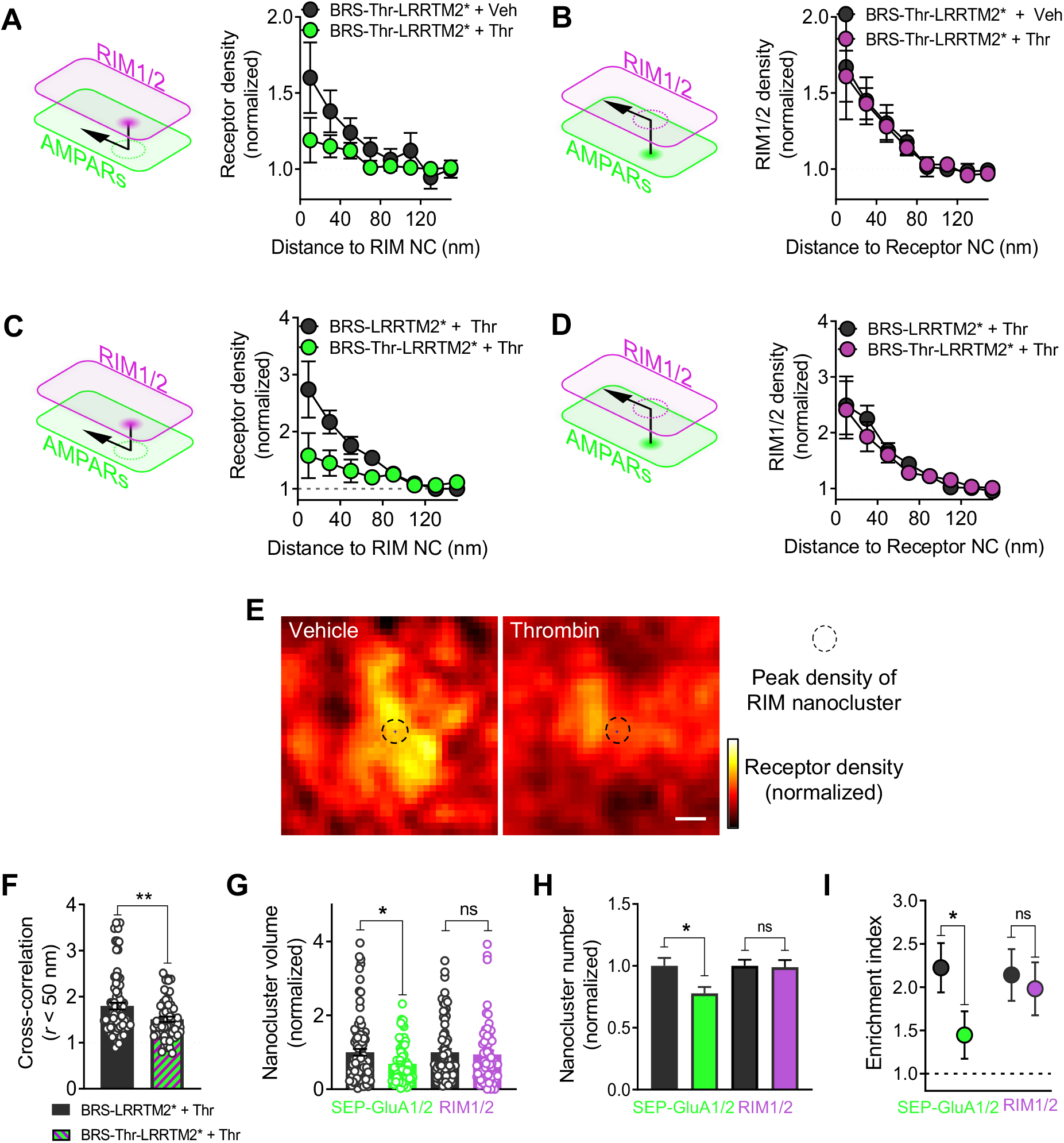
LRRTM2 is critical for AMPAR enrichment across from preferential sites of evoked neurotransmitter release. **(A)** Schematic demonstrating the measurement of AMPAR localization density across from RIM1/2 nanoclusters (left). Quantification of AMPAR enrichment from neurons co-expressing BRS-Thr-LRRTM2* and SEP-GluA1/2 following treatment with thrombin (10 minutes, green) or vehicle (black). **(B)** RIM1/2 density across from AMPAR nanoclusters from neurons in A. **(C)** Quantification of AMPAR enrichment from neurons co-expressing either BRS-Thr-LRRTM2* (cleavable, green; n = 95 nanoclusters/11 neurons/3 independent cultures) or BRS-LRRTM2* (non-cleavable, black; 127/15/3) and SEP-GluA1/2 following treatment with thrombin (10 minutes). **(D)** RIM1/2 density across from AMPAR nanoclusters as displayed in B. Quantification (cleavable, magenta; n = 90/11/3) of BRS-LRRTM2* (non-cleavable, black; n = 103/15/3) and AMPARs following treatment with thrombin (10 minutes). **(E)** Representation of AMPAR density across from RIM1/2 peak density averaged across many synapses. Scale bar: 50 nm. **(F)** Paired cross correlation g_c_(r) < 50 nm of synaptic protein pairs of SEP-GluA1/2 and RIM1/2 from thrombin treated neurons expressing BRS-LRRTM2* (black) or BRS-Thr-LRRTM2* (green). **(G)** Volume of AMPAR nanoclusters. **(H)** Number of detected AMPAR nanoclusters. **(I)** Enrichment indices (g*_r_* < 50 nm) for AMPARs across from RIM1/2 nanoclusters (grey, green, left) and RIM1/2 across from AMPAR nanoclusters (grey, magenta, right). Data are presented as mean ± SEM, * *p* ≤ 0.05, ** *p* ≤ 0.01, *** *p* ≤ 0.001. Mann Whitney rank-sum test was performed for F-I.

In these neurons, we applied either vehicle (aCSF) or thrombin to test the acute regulation of AMPAR organization by LRRTM2’s extracellular interactions. Remarkably, brief treatment with thrombin dramatically reduced the density of AMPARs directly across from RIM nanodomains (to 37.7 ± 9.1% of control enrichment index, Fig. 4A). Of note, RIM1/2 density across from detectable AMPAR nanodomains was unchanged (97.7 ± 12.2% of control, Fig. 4B), suggesting a strictly postsynaptic nanoscale re-organization of AMPAR patterning within the PSD, but also that the position of the detected AMPAR nanodomains relative to RIM1/2 nanoclusters were largely unchanged. Since we could not yet rule out that this was caused by specific or off-target effects of thrombin, we repeated the experiment except that neurons were transfected with either BRS-Thr-LRRTM2* or BRS-LRRTM2*, and both conditions received brief treatment with thrombin. Consistent with the prior result, thrombin treatment to neurons expressing the cleavable, but not the non-cleavable LRRTM2 resulted in a large reduction in AMPAR density across from RIM nanodomains (36.5 ± 27.3% of control enrichment index, Fig. 4C, Supplementary Fig. 7D), confirming that the effect is specific to the cleavage of the LRRTM2 ECD. Furthermore, RIM density across from these AMPAR nanodomains was again unaffected (92.5 ± 30.6% of control, Fig. 4D). To visualize this AMPAR de-enrichment another way, we calculated a 2D view of these data by aligning all receptor map data to the peak of their corresponding RIM1/2 nanocluster, producing a histogram of receptor density across from the RIM1/2 nanocluster peak averaged over all measured nanoclusters from many synapses (Fig. 4E). This output thus represents the average distribution of receptors arrayed in the synaptic membrane facing a vesicle that might fuse at the center of a RIM nanodomain. Following thrombin treatment, the peak of this receptor array is diminished, and receptors are dispersed so that they are much less concentrated directly in line with the RIM nanodomain. Together, the rapid effect of thrombin in these experiments demonstrates that LRRTM2 via its extracellular domain is actively involved in the nanoscale organization of AMPARs within established synapses.

Changes in either the density or arrangement of these AMPAR nanodomains could have produced changes to the enrichment measurements. To further discriminate how AMPAR organization changed upon LRRTM2 cleavage, we assayed a number of properties that these AMPAR nanoclusters exhibited. To assess the effect on RIM-AMPAR alignment without explicitly identifying nanoclusters of either protein, we measured the cross-correlation between AMPAR and RIM1/2 density distributions from the same neurons transfected with BRS-Thr-LRRTM2* or BRS-LRRTM2*. This measure showed a reduction following LRRTM2 cleavage (63.8 ± 1.3% of control, Fig. 4F), indicating that their relative density distributions became less similar, consistent with the above. Receptor nanoclusters were 67.5 ± 0.7% the volume of control receptor nanoclusters (Fig. 4G), and 28.3 ± 0.8% less numerous (Fig. 4H), while RIM1/2 nanocluster number and volume were not altered (94 ± 10.9%, 98 ± 5.7%, respectively; Fig. 4G,H). Furthermore, the summary enrichment data (an average of the data within 50 nm of the opposite nanocluster) demonstrated a significant decrease following LRRTM2 cleavage for AMPARs, but not RIM1/2 (Fig. 4I). Taken together, these data are consistent with a model of a postsynaptic nanodomain-specific de-enrichment of AMPARs near RIM nanodomains. These data provide strong evidence that while not acutely required for controlling AMPAR number in the PSD (Fig. 2D-F), LRRTM2’s ECD is critical for ongoing coordination of AMPAR density across from RIM nanodomains.

### Numerical model to predict effects of LRRTM2 loss on synaptic transmission

These effects offer a unique opportunity to explore how changes in the nanoscale spatial organization of AMPARs in the PSD could alter receptor activation and synaptic transmission. To address this theoretically, we predicted the magnitude of the potential effect using a model based on prior work. Prior models of glutamate release and diffusion along with receptor opening kinetics have established that AMPAR open probability decreases characteristically as their distance to the site of release increases (*50*, *51*). Beginning with this simplification enabled us to estimate the relative synaptic response after release given different receptor distributions within the PSD without explicitly simulating glutamate diffusion or receptor kinetics.

We generated simulated receptor maps based on several key synapse features obtained from our measurements and the literature (Fig. 5A,B and Supplemental Fig 6A). When subjected to our spatial analysis, this basal arrangement of simulated receptor positions recapitulated the autocorrelation (Supplementary Fig. 8B) and well reflected both the relative change in the enrichment profile (Supplementary Fig. 8C) and the relative change in the enrichment index (Supplementary Fig. 8D) for receptors as measured in our cultured neurons. To deduce how many AMPARs would need to leave the nanodomain to result in de-enrichment to the same degree as observed after cleavage of LRRTM2 (Fig. 4F, 4M), we removed different numbers receptors from the modeled nanodomain, placed them randomly within the PSD, and then compared the enrichment profile of the redistributed synapse to that of the original modeled synapse. A loss of ~16 AMPARs from the modeled nanodomain resulted in a ~60% reduction in density within the nanodomain and well recapitulated the experimentally observed decrease in enrichment (Supplementary Fig. 6C,D).

**Fig. 5.**
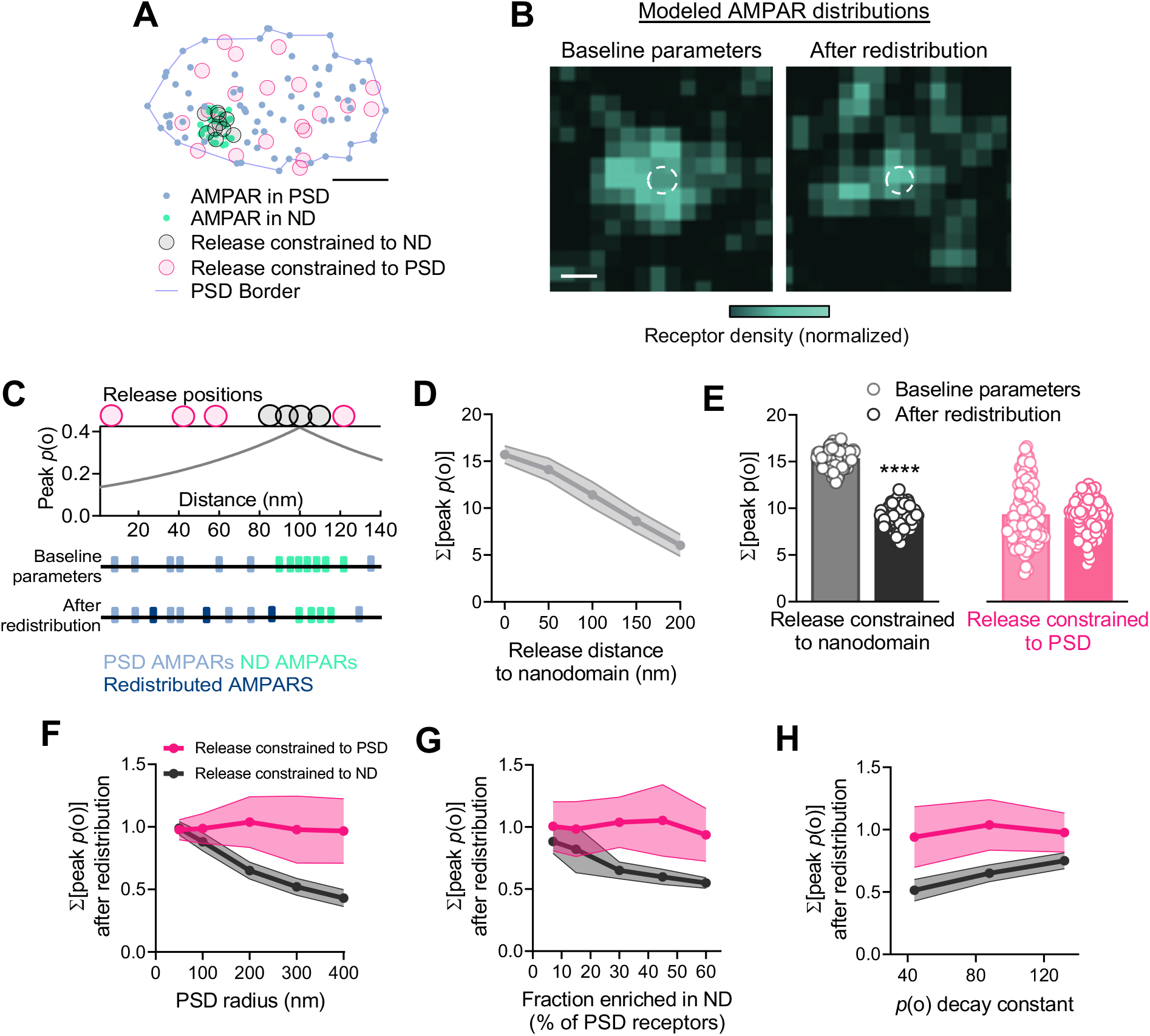
Numerical model to predict the effects of LRRTM2 loss on synapse function. **(A)** Example of the distribution of randomized AMPAR positions within the PSD and nanodomain and the distribution of randomized vesicle release positions, where release is constrained to either the boundary of the nanodomain (black) or PSD (magenta). Scale bar: 100 nm. **(B)** Representative density histograms of individual modeled receptor distributions. **(C)** Schematic demonstrating the calculation of peak open probability of all AMPARs within the PSD given a randomized release position constrained as described in a. **(D)** Calculation of the summed peak open probability of AMPARs as the release position is offset from the nanodomain. **(E)** Calculation of the summed peak open probability of AMPARs for release positions constrained to the nanodomain (left) and release positions constrained to the PSD (right) using the modeled nanodomain parameters (grey, black) and redistribution of AMPARs (pink, magenta). **(F)** Calculation of the summed peak open probability of AMPARs as PSD diameter is adjusted. **(G)** Calculation of the summed peak open probability of AMPARs as the proportion of AMPARs included in the nanodomain is adjusted. **(H)** Calculation of the summed peak open probability of AMPARs from as the peak open probability as a function of distance decay constant is adjusted ± 50%. Data in F-H are normalized to the baseline parameters condition. Data are presented as mean ± SEM. For all modeled data *n* = 100 randomizations/bin or 100 randomizations/condition. Mann Whitney rank-sum test was performed for E, **** *p* ≤ 0.0001.

Increasing evidence suggests that evoked and spontaneous transmission involve separable presynaptic structures (*52*) and activate separated groups of receptors (*53*), but their potential differential dependence on receptor nano-organization is not known. To model the impact of receptor redistribution on these different release modes, randomized release positions were constrained either to the nanodomain or the PSD as a whole to inform our predictions about evoked and spontaneous release, respectively (Fig. 5A, (*20*)). Then, by indexing the AMPAR peak open probability as function of distance from the vesicle fusion site (as in Tang et al., 2016, shown schematically in Fig. 5C, *20*), we calculated the mean summed peak open probability of all receptors in response to single glutamate release events before and after receptor redistribution. EPSC kinetics are not captured in such a model, but the mean summed peak open probability successfully indicated that release events closer to receptor nanodomains produce larger predicted responses (Fig. 5D) consistent with results observed from Haas et al. (2018; (*28*)).

To simulate the receptor reorganization observed after acute cleavage of LRRTM2, we removed the portion of the receptors from the nanodomain determined from the modeling above (16 of 27) and placed them randomly into the PSD outside the nanodomain. This manipulation substantially reduced the predicted response to release constrained to the receptor nanodomain (59.4 ± 0.1% of control; Fig. 5E, left). Thus, the model predicts that evoked transmission at “average” synapses would be reduced by roughly 60% after LRRTM2 cleavage. Strikingly, despite this strong effect, the response to release events occurring at randomized positions across the AZ was essentially unaltered (96.3% ± 0.1% of control; Fig. 5E, right). Interestingly, variability of response amplitude for spontaneous release was reduced upon redistribution (CV 33.03% for baseline parameters and 20.31% after redistribution), suggesting that heterogeneity of receptor density across the face of an individual PSD contributes to the response CV at that synapse.

This difference between the decrement of response to release at a nanodomain or across the synapse persisted over a range of parameters. For instance, we found that as PSD size increases, the proportional effect of receptor redistribution grows larger (Fig. 5F), suggesting that nano-alignment may be most critical for maximizing postsynaptic responsivity at large synapses. Similarly, positioning a greater fraction of receptors within the nanodomain resulted in a greater reduction in mean summed peak open probability upon redistribution (Fig. 5G) but again this only affected release constrained to the nanodomain, and no differences were observed in the mean of the responses with release constrained to the PSD.

A key parameter in the model is the decay profile in receptor open probability as a function of distance from the release site, here modeled by default as Po(d) = 0.42e-d/88 adapted from previous work (*20*, *50*). To test whether the outcomes were robust to changes in this parameter, we varied this decay constant by 50% in either direction. This altered the magnitude of influence as expected but did not qualitatively affect the outcome. When the decay rate was decreased (-d/44) making receptors less sensitive to release position, the mean summed peak open probability after nanodomain release events was elevated (not shown) yet was still strongly reduced upon redistribution (Fig. 5H). Conversely, when the rate of decay was increased (-d/132), responses were lower but also strongly reduced after redistribution. Note that for all values of the decay constant, in response to the modeled spontaneous release events, the mean summed peak open probability was essentially unchanged by redistribution of receptors (Fig. 5H).

Together, these simulations most generally suggest that receptor distribution within a PSD strongly influences the amplitude of evoked but not average spontaneous neurotransmission at that synapse. Specifically, they predict that following LRRTM2 cleavage, the amplitude of evoked EPSCs but not spontaneous mEPSCs should decrease substantially.

### LRRTM2 is critical for basal strength of evoked, but not spontaneous transmission

To assess these predictions of functional effects of AMPARs nanostructural remodeling within the PSD following LRRTM2 cleavage, we performed patch-clamp recordings from cultured hippocampal neurons while stimulating nearby cells to evoke synaptic responses (Fig. 6A, Supplementary Fig. 9A). In untransfected neurons, thrombin application had no effect on EPSC amplitude (98.8 ± 7.7% after 10 min, n = 9, Fig. 6B), confirming a lack of non-specific or endogenous effects of the protease. Similarly, in cells transfected with LRRTM2 knockdown and rescued with the non-cleavable GFP-LRRTM2*, EPSCs were unaffected (95.5 ± 5.8%, n = 9, Fig 6B). However, in cells transfected with GFP-Thr-LRRTM2*, acute cleavage of the LRRTM2 ECD resulted in a 45.3% ± 7.6% reduction in EPSC amplitude (n = 12, Fig. 6B), consistent with our modeling results.

**Fig. 6.**
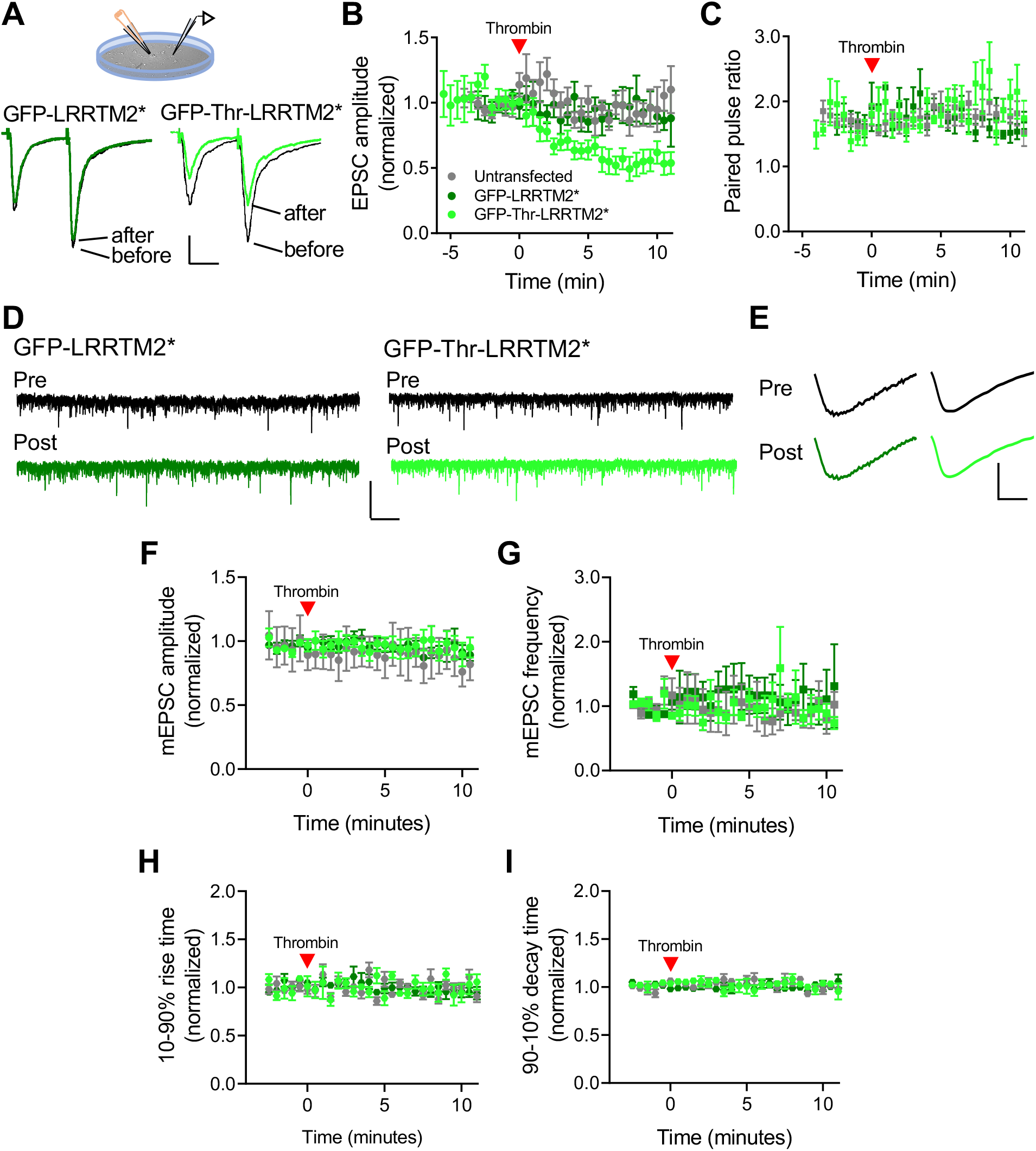
LRRTM2 is critical for basal strength of evoked but not spontaneous transmission. **(A)** Schematic of whole-cell patch clamp recordings from cultured hippocampal neurons with bipolar electrode stimulation to evoke synaptic currents. Below, averaged traces of evoked synaptic events. Neurons transfected with GFP-Thr-LRRTM2* (light green, cleavable, post-thrombin), GFP-LRRTM2* (dark green, non-cleavable, post-thrombin), where black indicates their respective baselines. Scale bar: 100 pA, 20 ms. **(B)** Quantification of evoked synaptic current amplitudes normalized to their baseline measurements over time from neurons expressing GFP-LRRTM2* (dark green; n = 9 neurons/3 independent cultures), GFP-Thr-LRRTM2* (light green; n = 12/3), or untransfected neurons (grey; n = 9/3). **(C)** Quantification of the paired-pulse ratio. **(D)** Representative traces of miniature EPSC (mEPSC) recordings from cultured hippocampal neurons transfected with GFP-LRRTM2* (dark green) or GFP-Thr-LRRTM2*(light green) before and after the application of thrombin. Scale bar: 35 pA, 2 s. **(E)** Averaged traces of miniature synaptic currents. GFP-Thr-LRRTM2* (light green) and GFP-LRRTM2* (dark green). Scale bar: 5 pA, 2 ms. **(F)** Quantification of miniature synaptic current amplitudes from GFP-LRRTM2* (n = 8/3), GFP-Thr-LRRTM2* (n = 8/3), or untransfected neurons (n = 5/3) before and after the application of thrombin. **(G)** Quantification of miniature frequency before and after the application of thrombin. **(H)** Quantification of the 10-90% rise times of mEPSC events over time, before and after the application of thrombin. **(I)** Quantification of the 90-10% decay time of mEPSC events over time, before and after the application of thrombin. Data are presented as mean ± SEM.

Deficits in presynaptic release probability could have contributed to the decreased EPSC amplitude. To test this, we calculated the paired-pulse ratio of responses to stimuli 50 ms apart. Thrombin treatment had no effect on the paired-pulse ratio across GFP-Thr-LRRTM2*, GFP-LRRTM2*, and untransfected neurons (Fig. 6C), as well as compared to their own baselines. These data suggest changes in release probability did not drive the changes in the evoked response amplitude after LRRTM2 cleavage.

To test the effect of LRRTM2 cleavage on the postsynaptic response to spontaneous release of glutamate, we measured miniature EPSCs (mEPSCs; Fig 6D-I) from neurons expressing GFP-LRRTM2* or GFP-Thr-LRRTM2*. Thrombin application did not change mEPSC amplitude in cells transfected with non-cleavable GFP-LRRTM2*. Neurons transfected with cleavable GFP-Thr-LRRTM2* also showed no changes to mEPSC amplitude following thrombin application (98.5 ± 3.7% of control, Fig. 6F, Supplementary Fig. 9B), consistent with the model’s prediction. Nevertheless, the coefficient of variation of mEPSC amplitude was smaller after thrombin exposure in neurons expressing GFP-Thr-LRRTM2* vs GFP-LRRTM2* (Supplementary Fig 9G), as expected based on modeling (Fig 5E). Together, these results suggest that the number of synaptic AMPARs was not changed upon the acute loss of the LRRTM2 ECD, consistent with results in Fig. 2 that synaptic content of both surface AMPARs and PSD-95 was unchanged within 30 minutes following LRRTM2 cleavage.

To further test if loss of the LRRTM2 ECD alters presynaptic mechanisms, we quantified mEPSC frequency. Neurons transfected with the non-cleavable LRRTM2 showed no changes in frequency after 10 minutes of thrombin exposure (91.7 ± 19.4% of control, Fig. 6G, Supplementary Fig. 9C), consistent with the lack of nonspecific or endogenous protease effects. Interestingly, neurons transfected with GFP-Thr-LRRTM2* also showed no changes following 10 minutes of thrombin treatment (Fig. 6G), further strengthening the idea that the LRRTM2 extracellular domain does not acutely regulate presynaptic release probability.

It was previously shown that extracellular cleavage of neuroligin-1 (NL-1) reduced evoked EPSC amplitude without changing in mEPSC amplitude, and this was attributed to a reduction of presynaptic release probability (*38*). As NL-1 and LRRTM2 share presynaptic Nrx as a ligand, we examined whether NL-1 cleavage also resulted in reorganization of transsynaptic alignment. Interestingly, the acute cleavage of NL-1 did not change the relative nano-alignment of RIM and AMPARs (Supplementary Fig. 10). This further distinguishes the unique role of LRRTM2 in maintaining synapse nanoarchitecture.

Numerical models of glutamate diffusion and AMPAR kinetics have demonstrated that glutamate release away from AMPARs delays the opening of these channels by decreasing the number of immediately doubly-bound receptors, leaving many in a singly-bound state and slowing the rise of the EPSC as the concentration of glutamate in the cleft quickly equilibrates (*54*). Unfortunately, the measured variability in EPSC kinetics in our approach (which did not stimulate single presynaptic neurons for analysis) appeared to be dominated by variation in axonal conduction and other presynaptic factors, precluding interpretation of EPSC kinetics. However, we tested whether mEPSC kinetics were disrupted after LRRTM2 cleavage. We quantified both normalized and raw 10-90% rise time (Fig. 6H, Supplementary Fig. 9C) and 90-10% decay times (Fig. 6I, Supplementary Fig. 9D) of mEPSCs from neurons transfected with either GFP-LRRTM2* and GFP-Thr-LRRTM2* before and after the application of thrombin. Representative averaged traces (Fig. 6E) and group quantification, however, demonstrated no change in either the rise time (101.8 ± 8.9% of control, Fig. 6H) or the decay time (100 ± 4.2% of control, Fig. 6I). These results in combination with the raw and normalized miniature amplitudes (Supplementary Fig. 6F, Supplementary Fig. 7B), which are also sensitive to changes in single channel kinetics, suggest no population changes in AMPAR single-channel kinetics following the cleavage of the LRRTM2 ECD. These results are consistent with our numerical model of AMPAR position within the PSD (Fig. 5), where the average distance of AMPARs to the modeled spontaneous release positions across the AZ was unchanged upon AMPAR redistribution.

Taking these data together, we conclude that the LRRTM2 extracellular domain is required for close positioning of AMPARs to sites of evoked vesicle fusion, and that this distribution of receptors preferentially enhances evoked, but not spontaneous postsynaptic response amplitude.

## Discussion

We used acute proteolysis of the LRRTM2 extracellular domain to test the idea that transsynaptic interactions in the synaptic cleft control molecular organization and function of established synapses, independent of synapse formation and on rapid time scales. We found that acute cleavage of LRRTM2 quickly led to dispersal of its extracellular domain from synapses and prompted a strong reduction in the strength of evoked but not spontaneous synaptic transmission. Based on several lines of evidence, we conclude that this reduction in transmission arose from nanoscale redistribution of AMPARs within the synapse away from sites of glutamate release. First, LRRTM2 is concentrated in the synaptic nanocolumn, heavily enriched in nanoscale subdomains containing PSD-95 and AMPARs, and aligned transsynaptically with RIM nanodomains in the active zone. Second, after cleavage of LRRTM2, AMPARs became less densely enriched across from RIM nanodomains, even though total AMPAR content in synapses as measured by immunostaining or live imaging was unchanged for at least 30 minutes. Third, despite marked alteration of AMPAR distribution, the acute disruption of LRRTM2 did not grossly alter synapse number, structure, or molecular content, and notably the total synaptic content of PSD-95 also did not change within this period of interest. Fourth, presynaptic function was unaltered, as judged by fully intact spontaneous release and evoked paired-pulse response ratio. Finally, numerical modeling of AMPAR activation based on our nanostructural measurements of AMPAR position well predicted the degree of reduction in evoked transmission, while also providing a mechanism for the lack of effect on the amplitude of spontaneous EPSCs. Together, these findings provide support for a model in which the nanoscale patterning of AMPARs is dynamically controlled by interactions of LRRTM2 with cleft proteins enriched within the nanocolumn, and that this organization can dramatically enhance AMPAR activation during evoked synaptic transmission.

The close correspondence between our measurements and the predictions from numerical modeling provide experimental support for the longstanding notion that receptor distribution within synapses affects synaptic strength (*5*, *20*, *55*). Modeling indicates that the combination of the sharp decay of glutamate concentration away from the site of fusion with the rapid kinetics of AMPAR activation and desensitization necessitate that AMPARs positioned closest to the site of glutamate exocytosis contribute proportionally most to EPSCs (*8*, *50*). The quick relaxation of EPSC amplitude towards a reduced steady state that we observed following LRRTM cleavage indicates that this mechanism likely plays a role in maintaining basal synaptic strength. Mechanisms that maintain synaptic strength absent the active induction of plasticity are not clear, but have been postulated to involve adhesion systems (*56*). Taken most broadly, our findings suggest that mechanisms of synaptic maintenance may be divided into those which establish the molecular constituent list including AMPAR number, and those which facilitate appropriate nanostructural organization. It is interesting to consider whether synapses of limited molecular complexity may adopt a somewhat disorganized configuration “by default,” whereas the presence of LRRTM2 enables organization into a configuration of higher synaptic potency.

There are a number of means by which LRRTM2 may organize AMPARs. Single-cell knockout of LRRTM1 and 2 in young adult mice reduced AMPAR-mediated evoked transmission and destabilized AMPARs as measured by photoactivation without affecting synapse number, release probability, or NMDAR-mediated transmission (*35*, *36*), suggesting that in established synapses, LRRTMs may serve as anchors for AMPARs in the PSD. However, the role of LRRTM1 in this process appears limited, since LRRTM1 knockdown alone has no effects on evoked or spontaneous EPSC amplitudes, but substantial impact on mEPSC frequency, spine density, and synaptic vesicle distribution (*57*) its function may principally be limited to presynaptic roles following synaptogenesis. These findings strongly suggest that LRRTM2 plays a unique role for AMPAR retention. However, disruption of LRRTM2 did not lead to loss of AMPARs from the synapses within 30 to 60 minutes, even though it eventually produced large changes in AMPAR stability in spines. Thus, LRRTM2 alone may not be sufficient to fulfill the functional role of “slots” hypothesized to anchor AMPARs (*58*, *59*). Further, we cannot rule out that the transmembrane and intracellular domains of LRRTM2 that remain after thrombin cleavage may contribute to the synaptic retention of AMPARs. This would be intriguing intramolecular segregation of function within LRRTM2 for overall retention vs positioning of AMPARs. However, GPI-anchored LRRTM2 ECD fully rescues the deficit in LTP during conditional deletion of LRRTM1 and 2, suggesting a specific role for the LRRTM2 extracellular domain.

At the subsynaptic scale, the patterned distribution of AMPARs is generally thought to be stabilized by a combination of factors in spite of continuous receptor diffusion in the plasma membrane (*15*, *60*, *61*): a heterogenous affinity landscape in the synapse created by the distribution of direct AMPAR binding partners (*62*), and an array of steric obstacles which creates macromolecular crowding and hinders their motion within the dense synaptic environment (*49*, *63*). We suspect both these mechanisms may be involved in how LRRTM2 controls the AMPAR pattern. There is some evidence that the LRRTM2 ECD can interact directly with AMPARs (*33*, *36*) (but see (*64*)), and LRRTM2 through its interaction with PSD-95 might dynamically organize intracellular scaffolds (*32*, *33*). At the same time, its loss may trigger reorganization or even loss of synapse-resident proteins which could alter the steric hindrance experienced by receptors in the cleft by their large extracellular domains or in the substantially denser PSD by their smaller intracellular domains. In addition, partitioning of the PSD via liquid-liquid phase separation is being actively investigated as a potential organizing mechanism of synaptic nanostructures (*65*). Due to multivalent interactions facilitated by LRRTM2 in the synaptic cleft, its presence could serve to establish a nanoscale, phase-separated synaptic subdomain into which AMPARs partition, and which would be disturbed by the cleavage of the LRRTM2 ECD (though we do not know of evidence that LRRTM2 is cleaved endogenously). Overall, regardless of the mechanism, our data indicate that the nanoscale organization of AMPARs is both modulated by the LRRTM2 ECD and capable of rapid reorganization.

These observations are particularly interesting given that strong evidence also implicates LRRTM2 in LTP (*36*). Conditional deletion of LRRTM1 and 2 in mature mice reduces LTP in vivo, and expression of the LRRTM2 ECD alone, but not LRRTM4, is sufficient to rescue these deficits (*35*), clearly consistent with our observation that acute loss of the ECD regulates synaptic strength. Similarly, point mutations to the LRRTM2 ECD which disrupt presynaptic neurexin binding fail to completely rescue LTP (*35*), suggesting that neurexin binding may help explain how LRRTM2 specifically organizes AMPARs with respect to active zone nanodomains. LRRTM2 has been proposed as an anchor that stabilizes AMPARs during LTP induction (*35*, *36*). Our work extends this by implying synaptic nanostructure shaped by LRRTM2 may play several roles in functional plasticity. Most simply, LRRTM2-augmented AMPAR activation may lower the threshold of activity needed to trigger plasticity. Existing LRRTM2 nanodomains could also facilitate the stabilization of recently exocytosed or otherwise labile receptors (*3*). Similarly, it is conceivable that LRRTM2 could nucleate new nanoclusters added during LTP (*66*, *67*), though activity-dependent trafficking of LRRTM2 remains uninvestigated. In addition, our prior observation that chemical LTP induction “sharpens” the AMPAR distribution under RIM nanodomains (*20*) may suggest further that graded levels of AMPAR organizational tuning could be facilitated by LRRTM2. Most broadly, it is a natural extension of our findings here to suggest that behavioral or disease-relevant plasticity mechanisms, even regardless of the potential involvement of LRRTM2, may regulate synaptic strength not only by regulating AMPAR number but through controlling synapse nanostructure.

Surprisingly, we found the average postsynaptic response to spontaneous release was rather insensitive to AMPAR nano-organization, though it remains to be seen if this holds for all synapse geometries (e.g. small synapses). Our model and others predict that different forms of release may produce different postsynaptic responses depending on the subsynaptic distribution of release sites. There is indeed evidence that mEPSCs as well as univesicular EPSCs evoked in the presence of Sr^2+^ differ in CV and amplitude from AP-evoked EPSCs in the same neurons (*68*, *69*), and our findings provide novel experimental support for the idea. However, one shortcoming in previous work as well as our own is that due to the large variation in synaptic potency even on single neurons, precise measures of both mEPSCs and evoked quantal size from the same *synapses* not merely the same cells will be needed for thorough experimental validation of these predictions. Such differences may be important though, because while the roles of spontaneous synaptic transmission remain unclear, it has been suggested to stabilize the basal structure and function of the postsynapse (*52*, *70*, *71*). Local activity driven by spontaneous neurotransmitter release is also important for restricting the lateral mobility of AMPARs, helping to trap them in the PSD (*72*). Implicating transcellular mechanisms in the distinct regulation of evoked and spontaneous transmission further distinguishes these two forms of transmission already known to operate with heterogeneity at different active zones (*73*).

The effects of disrupting LRRTM2 and NL-1 differ in several ways both electrophysiologically and molecularly. Perhaps most dramatically, our experiments showed that acute manipulation of LRRTM2 but not NL-1 quickly prompted disorganization of AMPARs, whereas in similar experiments, proteolytic cleavage of NL-1 rapidly altered synaptic neurexin content and reduced presynaptic release probability (*38*). Thus, even adhesion molecules that share binding partners may play unexpectedly divergent roles in maintaining organization of synaptic molecular complexes. More broadly, these results suggest that many specific aspects of synapse structure and function are maintained by unique subsets of the diverse cell adhesion systems present within single synaptic clefts. Indeed, growing evidence demonstrates that synaptic CAMs themselves are found in distinct subsynaptic patterns (*5*, *37*, *46*, *48*). Clearer understanding of CAM organization within synapses will provide insight into their contribution to synapse nanoarchitecture and to their cooperative or even competitive functional roles.

## Materials and Methods

### Plasmids

All LRRTM2 plasmids were generated based on FCK-shLRRTM2 and pBOS-GFP-hLRRTM2-FL described previously (*33*). For insertion of the thrombin cleavage site, the sequence coding the four Ser’s (S386-S389) was replaced with a sequence coding the cleavage site (LVPRGS) with a flexible linker (GGGGS) on each side. For knockdown-rescue experiments in neurons, the H1 promoter and sh-LRRTM2 sequences from FCK-shLRRTM2 were subcloned into the pBOS-GFP-hLRRTM2-FL around the MluI site with IVA cloning (*74*). For BRS-LRRTM2, GFP sequence was replaced with a sequence coding the α-bungarotoxin-binding sequence (WRYYESSLEPYPD; (*44*)). GFP-Neuroligin1 and GFP-Neuroligin1-Thr were kind gifts from Michael Ehlers (*38*).

### Co-culture synaptogenesis assay

Co-culture assays were performed as described (*75*). Briefly, neurons were dissociated from embryonic day 18 Sprague-Dawley rat hippocampi, plated at a density of 60,000 cells on 12 mm cover glasses pre-coated with 1 mg/ml poly-l-lysine (Sigma, P1274), and treated at 2 div for 24 with Ara-C (2 μM) to prevent glial growth. Neurons were cultured in Neurobasal medium (Invitrogen 21103-049) with 3% B27 (Invitrogen, 17504-001) and 1% Glutamax (Invitrogen, 35050-061) and incubated at 37°C in 5% CO2. When neurons reached 7 div, 70% confluent HEK293 cells were transfected using polyethylenimine in 6 well dishes at approximately 0.4 picomol plasmid per 9.5 cm2 well (*76*). Transfected HEK293 cells were suspended 24 h later and seeded onto 8 div neurons at a density of 5,000 HEK cells per 12 mm cover glass. Ara-C was added at 2 μM upon seeding to prevent HEK cell overgrowth. After 48 h, co-cultures were fixed on 10 div with 4% PFA, 4% sucrose in PBS, stained with primary antibodies diluted in 3% fetal bovine serum (FBS) and 0.01% Triton-X 100 in PBS, and incubated overnight at 4°C. Secondary antibodies were diluted in 3% FBS and applied for 4 hours at 4°C. Neuronal cultures were stained with mouse monoclonal antibodies against Bassoon (AssayDesigns Cat# VAM-PS003F, RRID: AB_2313991; 1:500) and BTX-Alexa-647. Secondary immunostaining was performed with Alexa dye-conjugated antibodies. Coverslips were mounted with Aqua/PolyMount (Fisher, NC9439247). Confocal microscopy was performed on a Leica TCS SP8. Images were acquired with an ACS APO 63x oil lens with 1.3 NA, using the same settings for each condition. During image acquisition and analysis, the researcher was blind to the condition. Images were analyzed using a custom written ImageJ script available upon request.

### Hippocampal Culture and Transfections

All experimental protocols were approved by the University of Maryland School of Medicine Institutional Animal Care and Use Committee or the Institutional Animal Care and Use Committees at the University of Science and Technology of China (USTC) and the Chinese Academy of Sciences (CAS). Dissociated hippocampal neurons from E18 SD rats of both sexes were prepared as described previously (*77*). Neurons were transfected on DIV 7-10 with Lipofectamine 2000 and experiments were performed at least 7 days after the transfection (DIV 14–21). All experiments were repeated on 3 or more separate cultures unless otherwise specified.

### Immunocytochemistry

Neurons were fixed in 4% paraformaldehyde, 4% sucrose in phosphate buffered saline (PBS) for 10 min at room temperature and processed for immunofluorescence with standard procedures as described previously (*20*). Primary antibodies were: rabbit anti-RIM1/2 (Synaptic System #140203, 1:500), mouse anti-PSD-95 (NeuroMab clone K28/43, 1:200), chicken anti-GFP (Chemicon ab13970, 1:200). Secondary antibodies were from Jackson ImmunoResearch (West Grove, PA), either already conjugated with Alexa 647 or unconjugated that we labelled with Cy3b (GE Healthcare). Labeling with the anti-GFP antibody was performed after fixation but prior to overt permeabilization.

Live cell α-bungarotoxin (BTX) recognition sequence (BRS) labeling with α-bungarotoxin conjugated to Alexa-647 was performed prior to fixation described above. Coverslips were inverted on 50 μl droplets of BTX-Alexa-647 (Thermofisher B35450, 1:100) in aCSF containing 2 mM Ca^2+^ and 2 mM Mg^2+^ and covered for 5 minutes at room temperature (21-24°C). Then, coverslips would be placed into a small weigh boat filled with aCSF, and gently agitated, removing and replacing the aCSF twice before mounting in the microscope imaging chamber.

For LRRTM2 and PSD-95 immunocytochemistry, neurons were fixed in 2% paraformaldehyde, 4% sucrose in cytoskeleton buffer (10 mM MES pH 6.8, 138 mM KCl, 3 mM MgCl_2_, 2 mM EGTA, 320 mM sucrose) for 8 minutes at room temperature. Coverslips were then washed 3 times for 5 minutes each with PBS/Gly. Cells were permeabilized with 0.3% TritonX-100 (TX-100) in PBS/Gly for 20 minutes at room temperature, then washed once in PBS/Gly with 0.1% TX-100 for 5 minutes. Blocking was performed with a solution containing 3% BSA, 5% goat serum, 5% donkey serum, and 0.1% TX-100 for 1 hour and 15 minutes. Coverslips were inverted and incubated with primary antibodies (αLRRTM2, IgG1A mouse, NeuroMab N209C/35.3, 1:10; αPSD-95, IgG2A, 1:80, stored in 50% glycerol) diluted in a 1:1 dilution of the blocking media and PBS/Gly overnight in a humidity chamber at 4°C. Coverslips were then washed 3x in PBS/Gly containing 0.1% TX-100 for 5 minutes. Secondaries (GαM IgG1A Alexa-647, Jackson, 115-605-205, Lot 143997, 1:200; DαM IgG2A, Jackson, Cat 20257, Lot 14C0225 1:200) were diluted in a 1:1 dilution of the blocking media and PBS/Gly and coverslips were inverted on secondary in a humidity chamber at room temperature for 1 hour. Then coverslips were washed 3x with PBS/Gly for 5 minutes. Cells were postfixed with 4% PFA, 4% sucrose in PBS for 15 minutes, then washed 3x with PBS/Gly for 5 minutes.

All imaging except for dSTORM and HEK co-culture assay was performed on an Andor Dragonfly spinning disk confocal on either an Olympus IX81 or a Nikon Ti2 microscope. In each case, a 60x/1.45 NA oil immersion objective and Zyla sCMOS camera were utilized, with image format of 103 nm/pixel. Except where indicated, experiments were conducted at room temperature (22-24°C). Otherwise, for imaging in culture medium, a stage-top incubator and objective heater (Tokai Hit) maintained the sample temperature at 37°C and CO2 at 5%.

### Proteolytic cleavage

Thrombin from bovine plasma (Sigma, T4648-1KU, Lot # SLBV3604) was diluted in the imaging buffer (aCSF; 2 mM Ca^2+^, 2 mM Mg^2+^) at 100 Units per mL such that when added to the bath by pipette the final concentration became 10 Units per mL. Thrombin was added drop-wise away from the objective into the media containing cells in the imaging chamber at 24°C. Thrombin was stored at −20°C with a volume of at least 600 μl, and only underwent 1 freeze-thaw cycle. For imaging at 0.003 Hz, cells were maintained at 24°C. Z-stacks were taken every 5 minutes, and a maximum intensity projection was used for analysis. For imaging at 20 Hz, cells were kept on the objective at 24°C. Imaging was performed with 488 nm excitation during continuous acquisition at 20 Hz. Binning (2×2) permitted the identification of modestly expressing synaptic puncta at lowest possible laser power to prevent photobleaching. Exposure was 50 ms per frame. Data was smoothed with a sliding average window with a bin length of 3 frames (150 ms). For all experiments, chamber was thoroughly washed with deionized water and 70% ethanol accompanied by physical scrubbing in order to completely remove residual thrombin that can adhere to the plastics and rubber of the imaging chamber and O-ring. All synaptic ROI measurements were background subtracted and normalized to an average of each synapses own baseline.

### Quantification of protein retention at synapses

#### Live (30 minute)

Cells were co-transfected with GFP-Thr-LRRTM2* or BRS-Thr-LRRTM2* and PSD95-mCherry* or SEP-GluA1,2, respectively. For PSD-95 experiments, multiposition z-stacks were acquired every five min. For analysis, maximum intensity projections were calculated. ROIs of a fixed size (15 pixels) were drawn around synaptic puncta containing both LRRTM2 and PSD95-mCherry* or SEP-GluA1/2 fluorescence. Integrated intensity was measured, background subtracted (an average of multiple ROIs across the field of view), then normalized to an average of the ROIs pre-thrombin baseline. Only spines that remained within the ROI for the duration of imaging were included.

#### Live (2 hour)

Cells were co-transfected with SEP-GluA1,2 and BRS-LRRTM2* or BRS-Thr-LRRTM2*. Multiposition z-stacks were taken every 15 or 20 minutes. For analysis, maximum intensity projections were calculated. ROIs of a fixed size (25 pixels) were drawn around synaptic puncta containing both LRRTM2 and SEP-GluA1,2 fluorescence. Integrated intensity was measured, background subtracted (an average of multiple ROIs across the field of view), then normalized to an average of the ROIs pre-thrombin baseline. Only spines that remained within the ROI for the duration of imaging were included.

#### Fixed

Cells were co-transfected with GFP-Thr-LRRTM2* and mCerulean3. Cells were immunostained for PSD-95 and RIM1/2 as described above. All regions were acquired with the same imaging parameters on the Dragonfly confocal. For analysis, background subtracted (values taken from an average of multiple background regions across the field) integrated intensity within an ROI of a fixed size (15 pixels). Values were additionally normalized to the median intensity in the field which helped to normalize potential differences in any region to region variability in staining intensity. Normalization to median intensity did not appear to be skewed by artefactual puncta as these were avoided during acquisition or occupied a very small fraction of total pixels in the field.

#### 24-hour post-thrombin

Cells were co-transfected with SEP-GluA1,2 and BRS-LRRTM2* or BRS-Thr-LRRTM2*. Both groups were treated with thrombin for 10 minutes and then returned to culture media. Then, 24 hours later cells were fixed and stained for GFP and the BRS-tagged LRRTM2 ECD as described above. All regions were acquired with the same imaging parameters on the Dragonfly confocal spinning-disk (Andor). For analysis, maximum intensity projections were calculated. ROIs of a fixed size (15 pixels) were drawn around αGFP puncta. Integrated intensity was measured, routinely background subtracted (an average of multiple ROIs across the field of view).

### Quantification of PSD-95 puncta density

Cells were transfected with cytosolic mCerulean3 alone or paired with either pBOS-shLRRTM2 (tgctattctactgcgactcde;(*33*)), GFP-Thr-LRRTM2* (this paper), or pBOS-GFP-Thr-LRRTM2 (this paper). Cells were then fixed and immunostained for PSD-95 (described above). mCerulean fluorescence was used to demarcate the dendrites of transfected cells. Using mCerulean fluorescence alone, in order to remain blinded to the transfection condition, up to the first 6 transfected cells were selected for imaging and further analysis in order to reduce bias. Regions were chosen at least ~75 μm from the soma when dealing with a clear primary dendrite to avoid volume effects. Distance was calculated by drawing a line in ImageJ (total pixel number x pixel size). All images were thresholded the same. Each punctum had to consist of at least 4 suprathreshold pixels. Experimenter was blind to the condition during image analysis.

### Quantification of spine morphology

Cells (DIV 4-6) were transfected with mCerulean3 alone or mCerulean3 paired with GFP-Thr-LRRTM2*. Cells were imaged at DIV 14-16. Maximum intensity projections of the confocal stacks were analyzed in ImageJ by an observer blinded to conditions. Analysis between groups were always performed within the same culture. For spine length, a line was drawn from the edge of the spine head to the edge of the dendrite, parallel to the long axis of the spine (total pixel number x pixel size). For spine area, an ROI was drawn around the spine head. The images were thresholded based on intensity and area was measured in ImageJ. Experimenter was blind to the condition during image analysis.

### Colocalization analysis

Cells were transfected with GFP-Thr-LRRTM2*. Synapses were picked based on colocalization with dendritic spines. Five consecutive spine-resident, GFP-positive puncta were selected randomly from at least 4 separate dendritic regions per cell when possible. When few branches were present, selection of dendritic regions of interest were as evenly distributed throughout the image as possible. The data represent the number of those randomly selected GFP-positive puncta which also contained at least 4 suprathreshold pixels of PSD-95 or RIM1/2 staining. Analysis was performed using ImageJ. Experimenter was blind during data analysis.

### 3D-STORM imaging

Imaging was performed essentially as described (*20*) on an Olympus IX81 ZDC2 inverted microscope with a 100×/1.49 TIRF oil-immersion objective. Excitation light was reflected to the sample via a 405/488/561/638 quad-band polychroic (Chroma) with an incident angle near but less than the critical angle. The typical incident power out of objective was ~30 mW for 647 nm and ~60 mW for 561 nm. Emission was passed through an adaptive optics device (MicAO, Imagine Optic) which corrected the aberrations and introduced astigmatism for 3D imaging. A Photometrics DV2 was insert before an iXon+ 897 EM-CCD camera (Andor) for simultaneous collection of the red and far-red emissions. All hardware was controlled via iQ software (Andor), except the MicAO which was controlled via Micromanager. Z stability was maintained by the Olympus ZDC2 feedback positioning system. Imaging of NL1 experiments was carried out on a Nikon ECLIPSE Ti2 inverted microscope equipped with a perfect focusing system and an 100×/1.49 TIRF oil-immersion objective controlled with NIS-Elements AR 4.30.02 software; emission was collected with a CMOS camera (ORCA-Flash4.0, Hamamatsu); localization detection, calibration and drift correction were done using the NIS-Elements AR analysis 4.40.00 software. Lateral drift was corrected with a cross-correlation drift correction approach24. Samples were imaged in a STORM imaging buffer freshly made before experiments containing 50 mM Tris, 10 mM NaCl, 10% glucose, 0.5mg/ml glucose oxidase (Sigma), 40 μg/ml catalase (Sigma), and 0.1M cysteamine (Sigma). TetraSpeck beads (100 nm; Invitrogen) immobilized within a thin layer of 4% agarose on a coverslip were localized across a z-stack with 30-nm steps to get the 3D calibration and correct alignment between the two channels as described previously. The average deviation of the bead localizations after correction was <15 nm in x/y directions and 40–50 nm in z.

### Single-molecule localization and analysis of synaptic clusters

All data analysis was performed offline using custom routines in MATLAB (Mathworks). The lateral (x, y) and axial (z) coordinates of single fluorophores were determined from the centroid position and ellipticity of the fitted elliptical 2D Gaussian function to a 7×7 pixel array (pixel size 160 nm) surrounding the peak. Poorly localized peaks were removed with a set of rejection criteria including an x–y precision <10 nm, fitting R2 > 0.6, and comprising >200 photons, and the shape of peaks24. For peaks lasting for more than one frames, only the localizations in the first frame were included in further analysis.

Synapses were identified as a juxtaposed pair of localization clusters of synaptic proteins and only those with clear pre- and postsynaptic components were selected for further analysis. A DB-SCAN filter was applied to the selected synaptic localizations with MATLAB function ‘DBSCAN.m’ created by S. M. K. Heris to define the boundaries of synaptic clusters. Only those localizations with a minimum of 60 localizations (MinPts = 60) within a radius of 5 times mean nearest neighboring distance (epsilon = 5 x MNND ≈ 100-120 nm) were considered within the synaptic cluster. The cluster boundaries were defined by an alpha-shape with α = 150 nm.

### Nanocluster detection and protein enrichment analysis

Nanoclusters within synaptic clusters were automatically identified based on local densities defined as the number of localizations within a certain distance (d) from each localization. To account for the variation in localization density across different synaptic clusters, we defined d as 2.5 _x_ MNND instead of a fixed value (*78*). The threshold of local density for nanocluster detection was defined as Mean(LD0) + 4 x Std(LD0), where LD0 is the local density of a randomized cluster with the same overall density as the synaptic cluster. The threshold we used represented the 99.95% confidence that the measured density differs from chance.

All localizations above the threshold were then ranked based on their local densities in a descending order and assigned each localization sequentially as the peak of a new nanocluster or a part of an existing nanocluster based on whether it was further enough from peaks of all existing nanoclusters. The localization with highest local density, if above the threshold, was defined as the peak of the first nanocluster. The second-highest-density localization would be considered as the peak of another potential nanocluster if the distance between the first peak to this potential second peak was larger than the defined cutoff distance; otherwise, the second localization was considered a part of the first nanocluster. The minimum peak-to-peak distance was set as 80 nm, which is about the average size of synaptic nanoclusters (*15*, *20*, *79*). Then, each potential nanocluster was further divided into sub-clusters based on the point-to-point distance with a cutoff of 2 x MNND using MATLAB function ‘clusterdata’, and only the sub-cluster having the original peak localization of this potential nanocluster was selected. Finally, the sub-clusters had to include at least 4 localizations to be accepted as a nanocluster.

The enrichment analysis is based on the prediction that if the pre- and postsynaptic nanoclusters align across the cleft, the presence of a nanocluster on one side will predict a higher local protein density around its projected point on the other side. The synaptic cluster pair was first translated to overlap with each other based on their general shape without bias towards local densities (*20*, *78*). The enrichment was then quantified as the average local density of protein A over the distance from the projected peak of a protein B nanocluster. In case of a positive alignment, this curve would start from a local density significantly higher than the average at the small distance and then decay to the average. More details and the defined MATLAB function for nanocluster detection and protein enrichment analysis could be found in Chen, et al., 2020. Experimenter was blind during image analysis.

### Automatic enface projection and averaging of synapses

A plane parallel to the cleft was defined by fitting all localizations after the translation (least square of the normal distance to the plane). The two-dimensional enface projection was achieved with calculation of the projected coordinates of all localizations along the fitted plane. To avoid the potential dilution of local density after the collapse of one dimension, maximal projection of 3D local density was made to generate the density map of projected cluster. To visualize the enface distribution of both RIM1/2 and PSD-95 around PSD-95 nanoclusters, we averaged both normalized density maps centered around the projected peaks of PSD-95 nanoclusters. Meanwhile, to avoid any artifact created by the bordering effect, all values outside the synaptic cluster were replaced with 1 before averaging was performed (*78*).

### Numerical model to estimate peak open probability of AMPARs

We used a constrained deterministic approach to test how different AMPAR organizations could impact the peak open probability of AMPARs at individual synapses based on key biological measurements. The calculation of ∑[peak open probability], also denoted as∑[peak p(o)], is adapted from previous stochastic modeling (*20*, *50*), where the probability of channel opening characteristically decays as a function of the distance to the position of glutamate exocytosis. This relationship has been modeled here as Po(r) = 0.42e-r/88, as described previously (*20*).

AMPAR positions were randomly generated in MATLAB using cirrdnPJ.m which creates points within a circle of a specified size, essentially building a map of randomized AMPAR positions with 2D coordinates. We considered this the modeled PSD area, and this area was determined by values taken from prior EM work (*80*). The number of points to be generated within the PSD area was taken from prior work (*81*). Using a separate loop of cirrdnPJ.m, another smaller radius could be specified within the larger PSD area, in which points were randomly generated. We considered this the modeled nanodomain, and it contained the average number of AMPARs suggested to form these nanodomains (*79*).

These modeled AMPAR organizations containing a single nanodomain were examined using our spatial analysis. The autocorrelation measurement, as described previously (*20*) for both biological and modeled localization data, was used to measure the size of these modeled subsynaptic clusters. The detected size of the modeled nanodomain was similar to the subsynaptic organizations observed in the biological data where the profile decays back to 1 at ~80 nm indicating the size of the modeled nanodomain. Of course, AMPAR nanodomains found in biological synapses can range in number impacting the amplitude of this measurement due to the increased frequency of the signal, and multiple nanodomains in one synapse will show a larger amplitude when measured by the autocorrelation. Since we only model a single nanodomain within the synapse, the autocorrelation correctly demonstrates a lower amplitude than the measured biological data.

These modeled AMPAR organizations were examined using our enrichment analysis (*20*, *78*). This analysis measures the density of points as a function of distance beginning at a specified position and moving out radially at determined step sizes (distance in nm). Using the previously described baseline parameters, and in agreement with the autocorrelation, the enrichment analysis successfully demonstrated that these modeled AMPAR positions show subsynaptic enrichment decaying to the average synaptic density by ~80 nm, and this measure was expectedly sensitive to the number of AMPARs included in the nanodomain. Then, the sensitivity of the threshold-based nanocluster detection algorithm, which can detect the number of points included in a subsynaptic cluster was adjusted until it successfully indicated that on average ~27 AMPARs were in-nanodomain. In order to reflect the redistribution of AMPARs within synapses observed in the biological data by dSTORM, some number of AMPARs had to be removed from the nanodomain, but not lost from the PSD (Fig. 2, Fig. 6), which is referred to here as ‘redistribution.’ The specific mechanisms driving AMPAR position after this redistribution remain unclear, for instance, whether AMPARs are specifically excluded following LRRTM2 cleavage has not been determined. To reflect this in our model, AMPARs were simply placed back randomly into the modeled PSD, thus not specifically excluded from the nanodomain area after redistribution. Then using the enrichment profile and enrichment index as readouts of this reorganization, AMPARs were redistributed using this approach until the modeled enrichment index and modeled enrichment profile closely approximated the difference in the measured enrichment index and measured enrichment profile of AMPARs in biological synapses compared to their respective controls. Then, using the nanocluster detection algorithm adjusted to detect the modeled ‘ground truth’ number of AMPARs previously, we redistributed AMPARs and quantified how many AMPARs were still considered to be ‘in nanodomain’ after redistribution. Interestingly, some nanodomains were no longer detectable given the magnitude of reorganization, which is consistent with our observations in Fig. 4l.

The open probability of AMPARs is thought to critically depend on the distance to the site of glutamate exocytosis. Release position has been thought to occur in spatially distinct subregions of the active zone given different modes of neurotransmitter release, thus influencing this key parameter. To understand how different constraints on release position in the active zone impact AMPAR open probability, we modeled two modes of release, again using cirrdnPJ.m to randomize AMPAR positions within a specified area. Release constrained to the nanodomain is referred to here as ‘evoked release,’ as it is thought to occur over a smaller fraction of the PSD and aligned with postsynaptic AMPAR nanodomains. We refer to release over the entire area of the PSD as ‘spontaneous release’, as it does not demonstrate a similar constraint in release position distribution determined by live-imaging of vesicle fusion events (*20*).

Then, using the baseline parameters as a starting point, and our modeled redistribution of AMPARs given the data from Fig. 4F,M (as described above), we calculated the open probability of each AMPAR in the modeled PSD. This was done by indexing the distance of each AMPAR to the modeled vesicle release position. Then by summing these probabilities across every AMPAR in the modeled PSD, we could estimate the number of AMPARs on average that would generally be expected to open in response to spontaneous or evoked release. This informed the interpretations of the electrophysiology from neurons that underwent LRRTM2 cleavage and subsequent AMPAR redistribution. To test how these various spatial parameters (‘PSD area’, ‘proportion of AMPARs in the nanodomain’, and the ‘decay constant of the decay profile’) used in the model influenced AMPAR open probability, we kept the baseline parameters constant except for the parameter being tested and repeated the redistribution of AMPAR positions described above over a reasonable biological range.

### Electrophysiology

Whole-cell recordings were made on neurons from DIV13-17 with 5-8 MΩ pipettes filled with an internal solution that contained (in mM): 130 CsMeSO_3_, 6 NaCl, 10 HEPES, 1 MgCl_2_, 2 BAPTA-K, 0.2 CaCl_2_, 3 Mg-ATP, 0.3 Tris-GTP, pH 7.3 with CsOH, 290-295 mOsm. Neurons were hold at −70 mV at which the GABAergic current were minimal. The bath solution consisted of (in mM): 130 NaCl, 2.5 KCl, 1 NaH_2_PO_4_, 10 HEPES, X CaCl_2_, 4-X MgSO_4_, and 10 Glucose. Lower [Ca^2+^]_o_ (X = 0.5-1) was used for eEPSC recordings to reduce the recurrent activity, while for mEPSC recordings normal [Ca^2+^]_o_ (X = 2) combined with TTX (1 μM) and picrotoxin (50 μM) was used. Evoked EPSCs were elicited with 1-ms extracellular field stimuli through a bipolar electrode made from θ-shape glass pipette with opening of 2 to 3 μm. The stimulation electrode was held a few μm above the cells and moved around to locate at a position where single-peak, monosynaptic currents were reliably evoked. The paired stimuli with a 50 ms interval were delivered every 10 s. For miniature EPSCs, glass pipettes were pulled to have a resistance of 3-6 MΩ. An internal solution containing 130 mM K-gluconate, 5mM KCl, 2 mM MgCl_6_-H2O, 10 mM HEPES, 4 mM Mg-ATP, 0.3 mM Na_2_-GTP, 10 mM Na_2_-phosphocreatine, and 1 mM EGTA was used to record at room temperature (21-24°C). The series resistances were monitored and data with changes of >20% were discarded. The capacitance and input resistance were not significantly different between different groups of neurons. Data were collected with MultiClamp 700B amplifiers (Molecular Devices) and digitized at 5 kHz with Digidata 1440 and Clampex 10 software (Molecular Devices). mEPSCs were detected by fitting to a variable amplitude template using pClamp 10 analysis software. Experimenter was blinded to condition during data analysis.

### Statistical analysis

We used two-tailed Student’s T or Mann-Whitney rank sum tests for comparisons between 2 groups. We used a one-way ANOVA with posthoc Dunnett’s Test for multiple comparisons of three groups. Data are presented as means ± SEM except otherwise noted. Significance levels displayed as follows: n.s., not significant, *p* > 0.05; * *p* < 0.05, ** *p* <0.01, *** *p* <0.001, **** *p* <0.0001. These tests were performed in Prism 8.2.0 (GraphPad).

## Acknowledgments

The authors thank Minerva Contreras for outstanding assistance and preparation of cell cultures and members of the Blanpied lab for their comments and help on experiments, analysis, and this manuscript.

## Funding

National Institute of Mental Health F31MH116583 (AMR)

Brain & Behavior Research Foundation NARSAD Young Investigator Award and National Natural Science Foundation of China 31872759 (AHT)

National Institude of Mental Health R01MH086828 (SMT)

National Institute on Drug Abuse R01DA018928 (TB)

National Institute of Mental Health R01MH119826 (TAB, TB)

National Institute of Mental Health R37MH080046 (TAB)

## Author contributions

A.H.T., A.M.R., T.B., and T.A.B. designed the experiments. A.M.R., A.H.T., T.A.L., X.Z.G, and B.A.C. conducted the experiments and data analysis. A.M.R., A.H.T., and X.Z.G performed the single-molecule imaging experiments. A.H.T., A.M.R., and T.A.L. performed the electrophysiology experiments. B.A.C. and T.B. performed and analyzed the co-culture experiments. A.M.R. drafted the original manuscript. A.M.R., A.H.T., and T.A.B wrote and revised the manuscript. All authors contributed to editing the manuscript and evaluating the data.

## Data and materials availability

Data and materials are available upon request.

