## Supplemental figures for "Subsynaptic positioning of AMPARs by LRRTM2 controls synaptic strength"

||Present address: Ohio State University College of Medicine, Columbus, OH, USA.

† These authors contributed equally
* Corresponding author

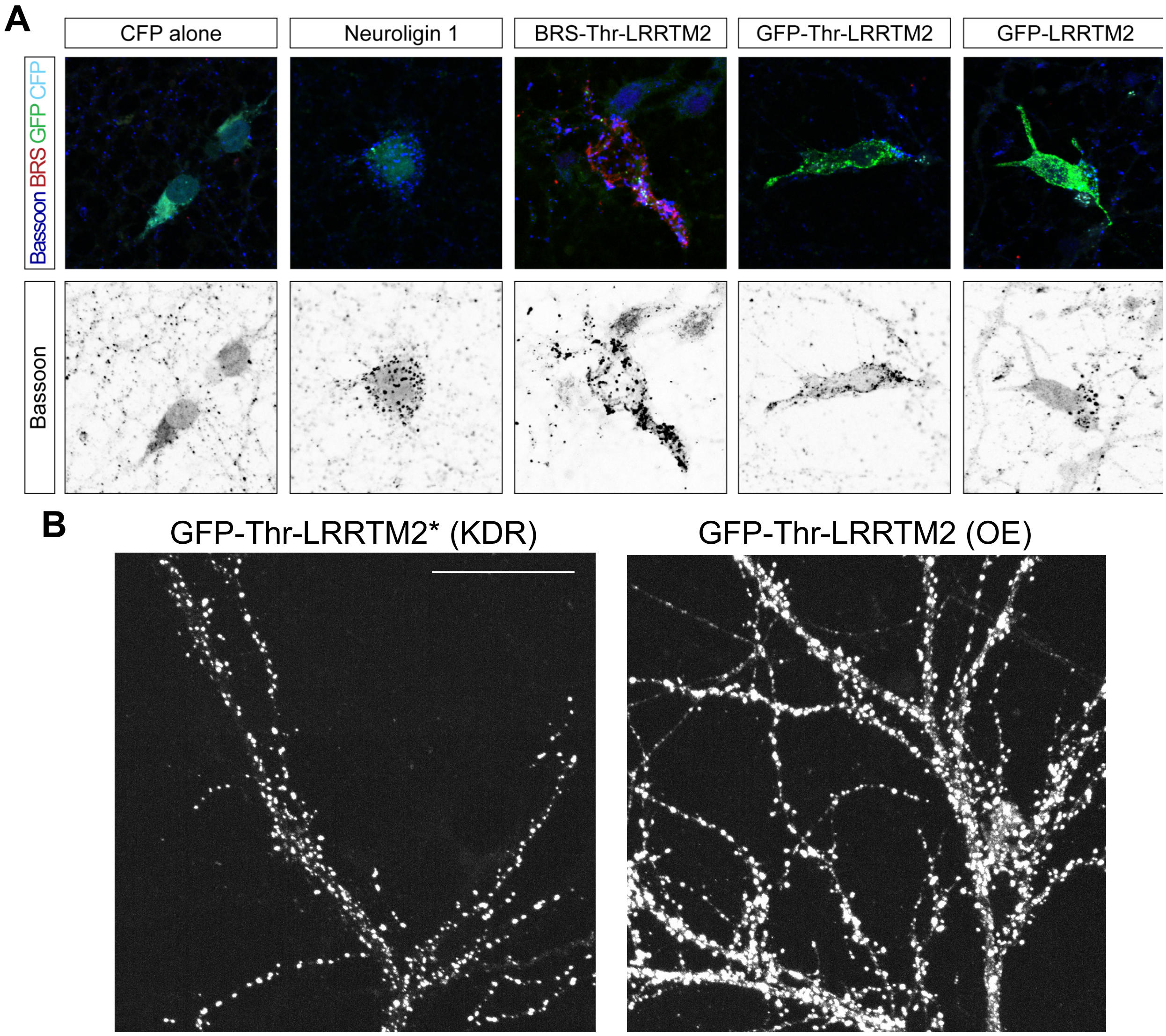


**Fig. S1.** Accompanies Fig 1. **(A)** Representative images from HEK-neuronal co-culture synaptogenesis assay. **(B)** Representative images from neurons expressing the knockdown-replacement vector GFP-Thr-LRRTM2* or overexpressing GFP-Thr-LRRTM2 demonstrating elevated synaptic enrichment of LRRTM2 in the knockdown-replacement. Scale bar: 20 µm.

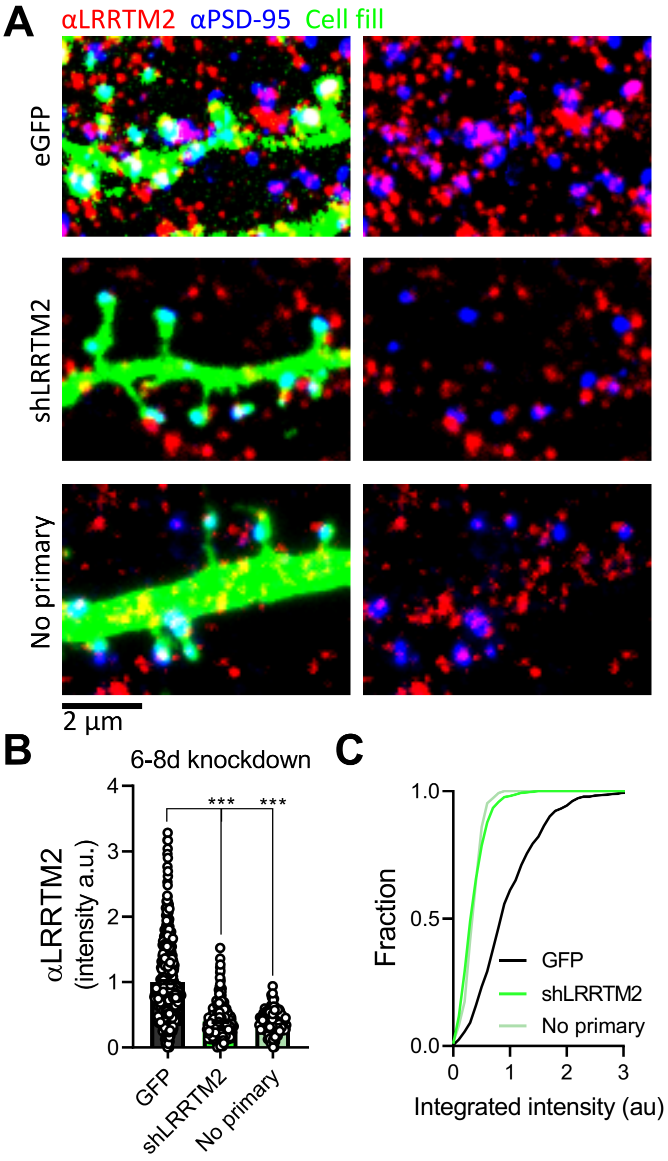


**Fig. S2.** Accompanies Fig 1. **(A)** Representative images from neurons transfected with cytosolic GFP (22 neurons/550 synapses/2 cultures), LRRTM2 shRNA (20/500/2), and staining lacking the primary antibody against LRRTM2 (175/7). **(B)** Quantification of LRRTM2 signal at PSD-95 positive spines normalized to control. **(C)** Cumulative distribution of LRRTM2 signal intensity Scale bar: 2 µm. Kruskal-Wallis with Dunn's multiple comparisons test, $p < 0.05$ = *, $p < 0.01$ = **, $p < 0.0001$ = ***.

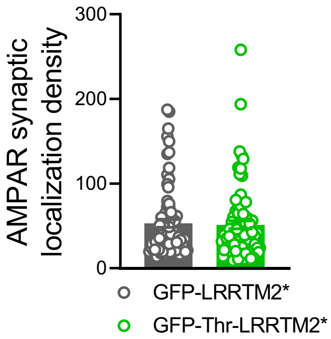


**Fig. S3. Synaptic localization density of AMPARs.** Accompanies Fig 2. Expression of GFP-LRRTM2* (grey) or GFP-Thr-LRRTM2* (green) in cultured hippocampal neurons and immunolabeling of SEP-GluA1/2 and RIM1/2 following 10-minute treatment with thrombin. Quantification of the synaptic localization density of AMPARs analyzed in Fig. 4c (non-cleavable, grey; 77/15/3; cleavable, green; n = 73/11/3).

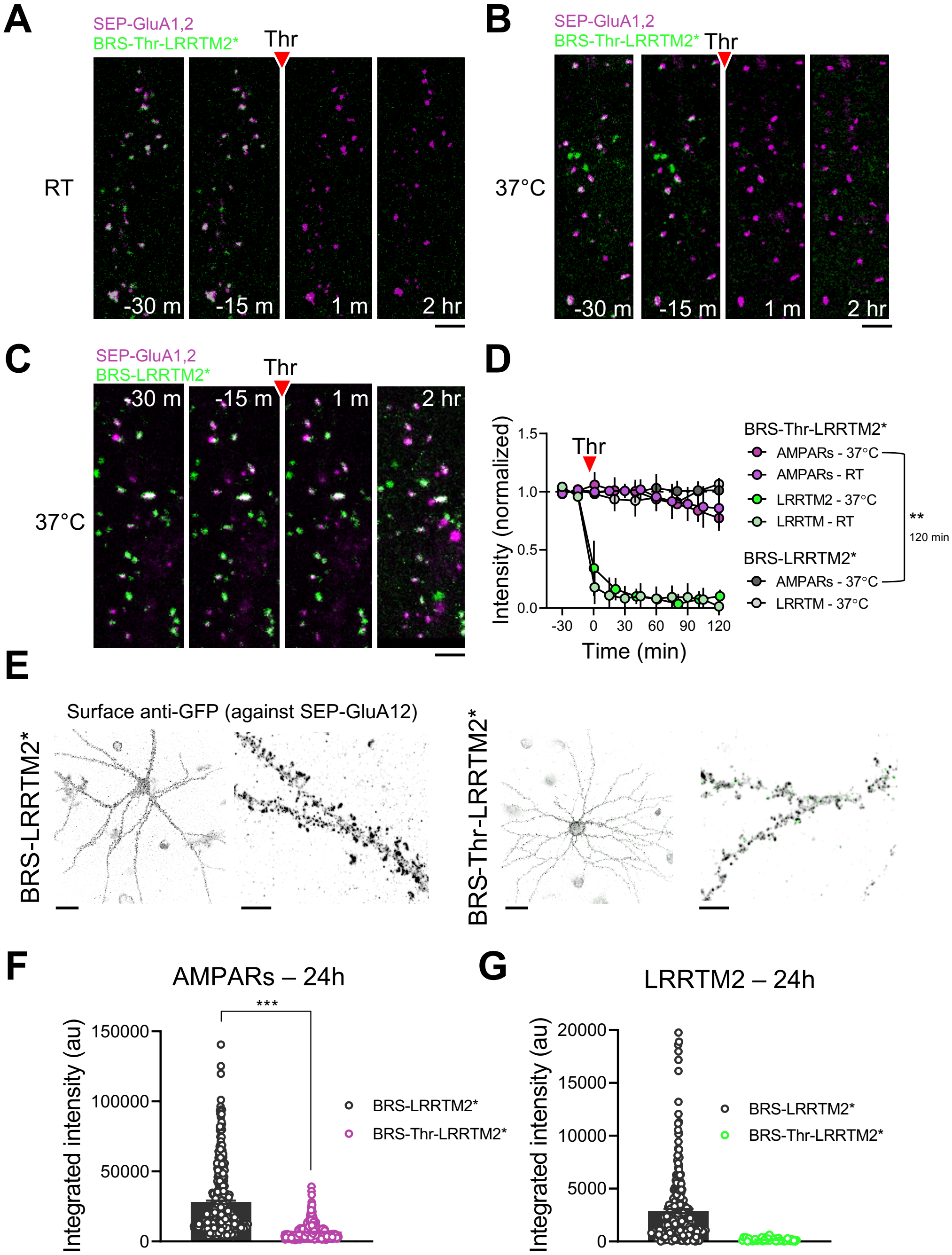


**Fig. S4.** Accompanies Fig 2. **(A)** Representative images from time series from neurons expressing SEP-GluA1,2 and BRS-Thr-LRRTM2* at room temperature (RT) in aCSF (n = 8 neurons/160 synapses/3 cultures), **(B)** SEP-GluA1,2 and BRS-Thr-LRRTM2* at 37°C in culture media (*n* = 6/120/2)*,* or **(C)** SEP-GluA1,2 and BRS-LRRTM2* at 37°C in culture media (*n* = 4/80/2). Scale bar: 2 µm. **(D)** Quantification from A-C. **(E)** Surface staining of SEP-GluA1,2 in cells expressing either BRS-LRRTM2* (*n* = 8 neurons/400 synapses/2 cultures) or BRS-Thr-LRRTM2* (11/550/2) 24 hours after treatment with thrombin for 10 minutes. Scale bar: 25 µm, 5 µm. **(F)** Quantification of SEP-GluA1,2 staining intensity. **(G)** Quantification of LRRTM2 labeling intensity. (D) Student's t-test, (F) Mann Whitney rank-sum test, $p < 0.05$ = *, $p < 0.01$ = **, $p < 0.0001$ = ***.

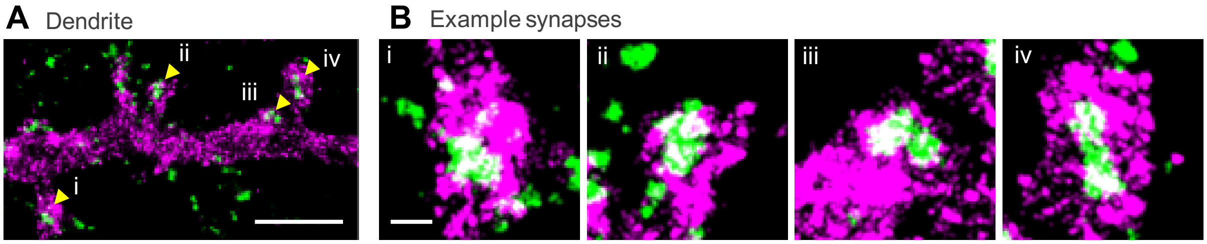


**Fig. S5.** Accompanies Figure 3. Representative images of SEP-TM distribution in dendrites as measured by 3D dSTORM, demonstrating a much broader distribution than the synaptic marker RIM. Scale bar a, 2 µm, b, 250 nm.

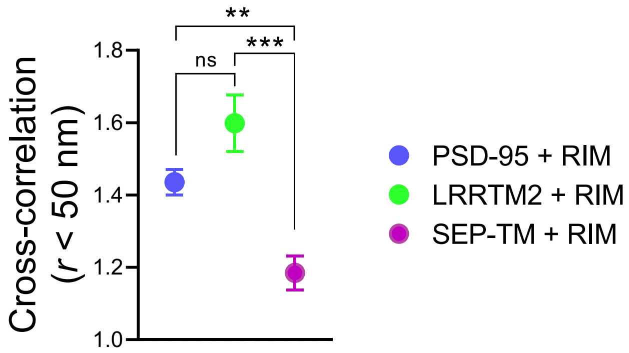


**Fig. S6.** Accompanies Fig. 3. Cross-correlation for LRRTM2 (n = 99 synapses/5 independent cultures), PSD-95 (n = 60/2), and SEP-TM (n = 66/3), where the correlation is performed between each molecule's cluster distribution and the synaptic distribution of RIM1/2. Kruskal-Wallis with Dunn's multiple comparisons test, $p < 0.05$ = *, $p < 0.01$ = **, $p < 0.0001$ = ***.

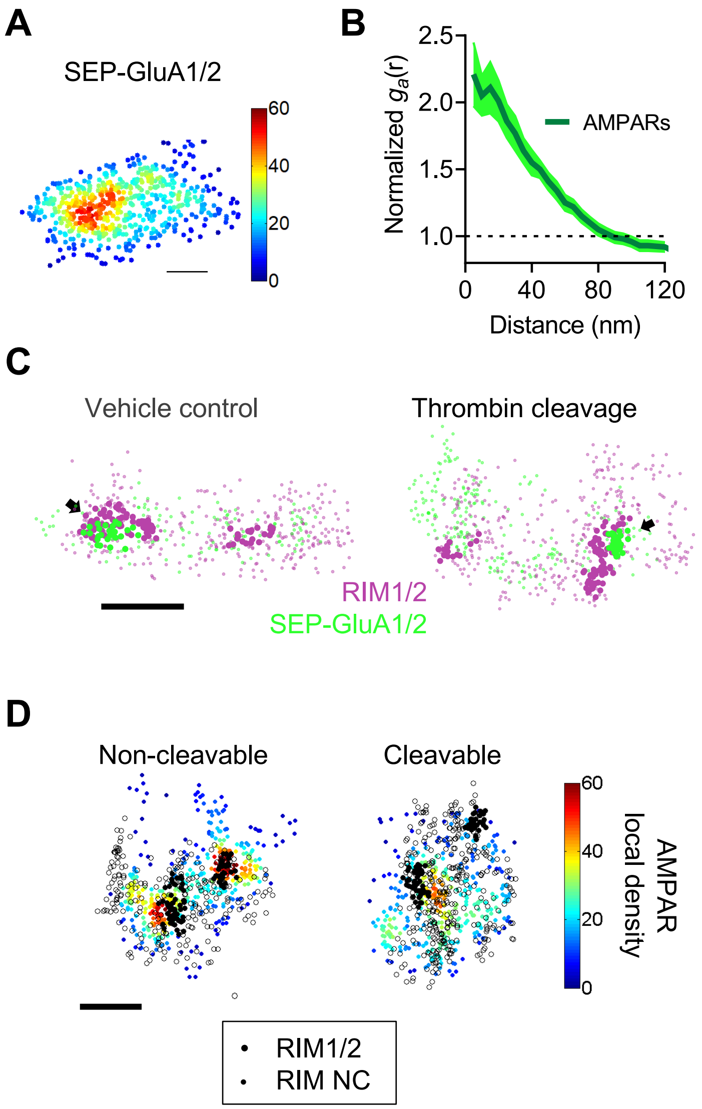


**Fig. S7.** Accompanies Fig. 4. **(A)** En face view of SEP-GluA1/2 localizations coded by their local density (5xNN distance). Scale bar: 80 nm. **(B)** Auto-correlation of SEP-GluA1/2, decaying to 1 by ~80 nm. (n = 55 synapses, 5 neurons). **(C)** En face views of synapses treated with either aCSF (left, vehicle control) or thrombin (right) labeled for RIM1/2 (magenta) and SEP-GluA1/2 (green) imaged with 3D dSTORM. Arrows and bolded localizations indicate detected nanoclusters. Scale bar: 100 nm. **(D)** En face views of synapses treated with thrombin while expressing either BRS-LRRTM2* (non-cleavable) or BRS-Thr-LRRTM2* (cleavable) imaged with 3D dSTORM. RIM localizations are denoted by the open black circles and filled black circles represent nanocluster localizations. SEP-GluA1/2 localizations are denoted by the jet lookup table, coded by their local density (5X nearest neighbor distances) where warmer colors indicate a higher local density. Scale bar: 100 nm.

**A**

Baseline parameters

| | | |
|---|---|---|
| PSD area | 0.12 µm$^2$ | Harris et al. (1989) |
| Nanodomain area | 0.005 µm$^2$ | MacGillavry et al. (2013) |
| Receptors in PSD | 100 | Matsuzaki et al. (2001) |
| Receptors in ND | 27 | Nair et al. (2013) |
| Peak *p*(open) | $0.42e^{-d/88}$ | Raghavachari et al. (2004) |
| | | Tang et al. (2016) |
| De-enrichment | 62% | *This paper* |

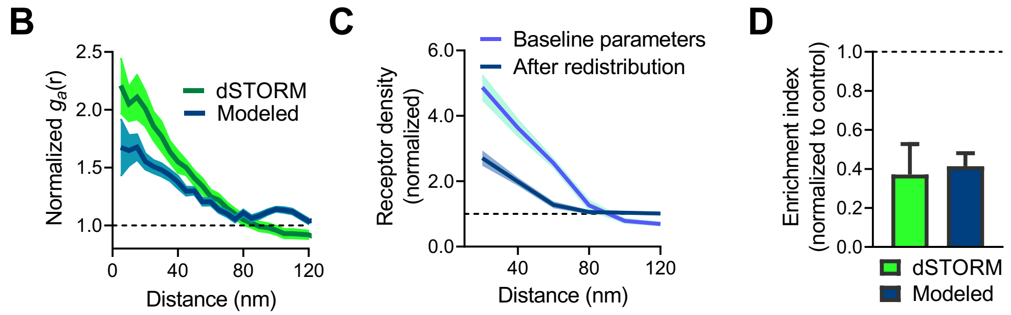


**Fig. S8.** Accompanies Fig. 5. **(A)** Baseline parameters incorporated into the model to estimate AMPAR peak open probability from modeled receptor organizations and release site positions. **(B)** Auto-correlation of measured SEP-GluA1,2 (from experiments in Fig. 4b, n = 55 synapses, 5 neurons) and modeled AMPAR organizations with a single modeled nanodomain (n = 100 randomizations). **(C)** Distribution of AMPAR density as a function of the distance to the modeled nanodomain normalized to the average synaptic density and the impact upon this self-enrichment following the redistribution of AMPARs from within the nanodomain to out of the nanodomain. **(D)** Normalized enrichment indices (gr(<50 nm)) for measured SEP-GluA1/2 (Fig. 4m) and modeled AMPARs following LRRTM2 cleavage and AMPAR redistribution, respectively.

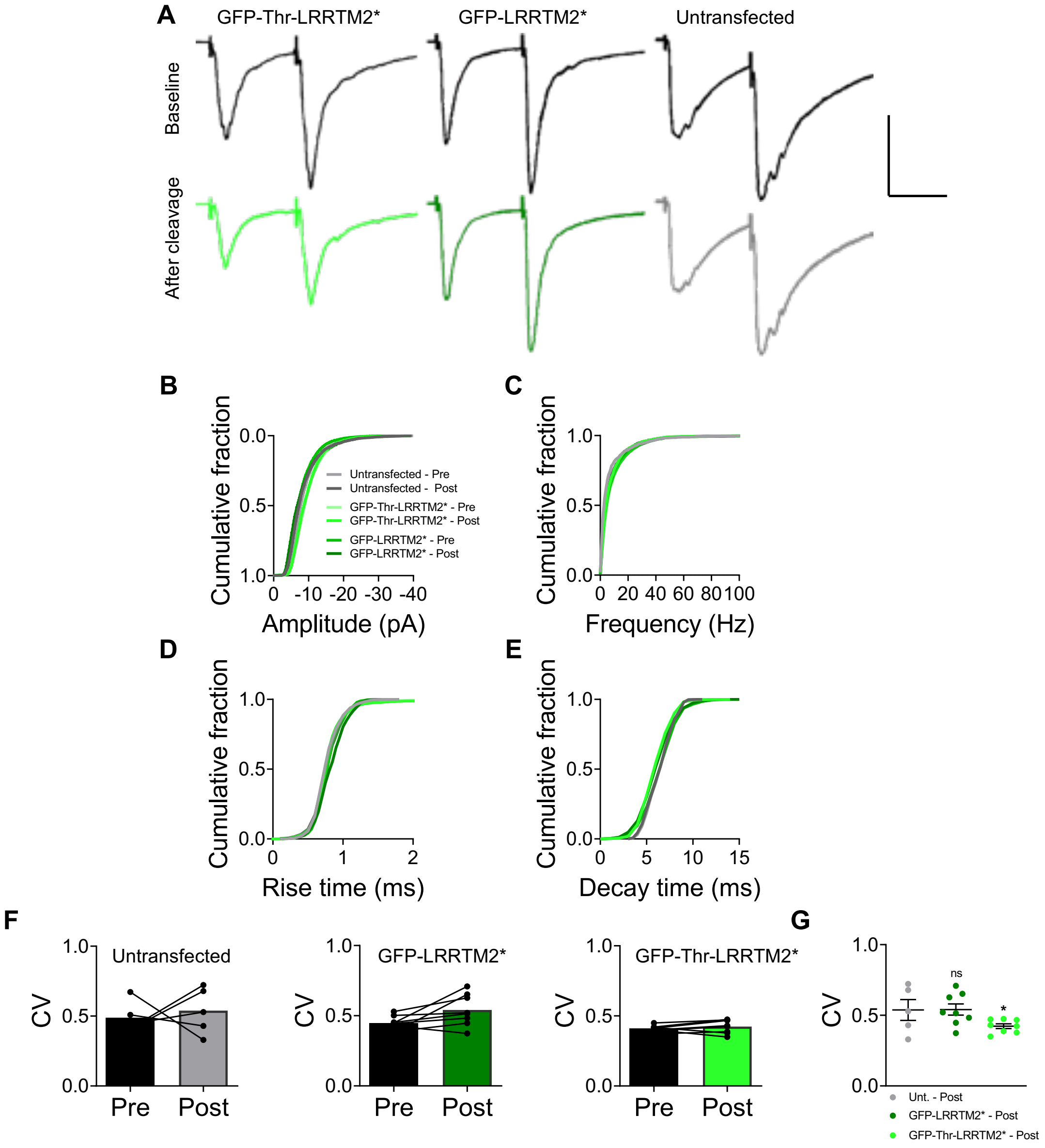


**Fig. S9. Synaptic responses before and after the application of thrombin.** Accompanies Fig 6. **(A)** Traces from Fig 6a, replotted offset from one another for clarity. Scale bar, 100 pA, 20 ms. **(B-G)** Quantification of spontaneous mEPSCs from neurons transfected with GFP-LRRTM2* (n = 8 neurons/3 independent cultures), GFP-Thr-LRRTM2* (n = 8/3), or untransfected neurons (n = 5/3) before and after the application of thrombin. **(B)** Amplitude, **(C)** frequency, **(D)** 10-90% rise time, **(E)** 90-10% decay time, **(F)** and coefficient of variation (CV) for each neuron for pre- and post-thrombin bins. **(G)** Summary of post-thrombin CV among these groups. One-way ANOVA with Dunnett's multiple comparison, $p < 0.05$ = *.

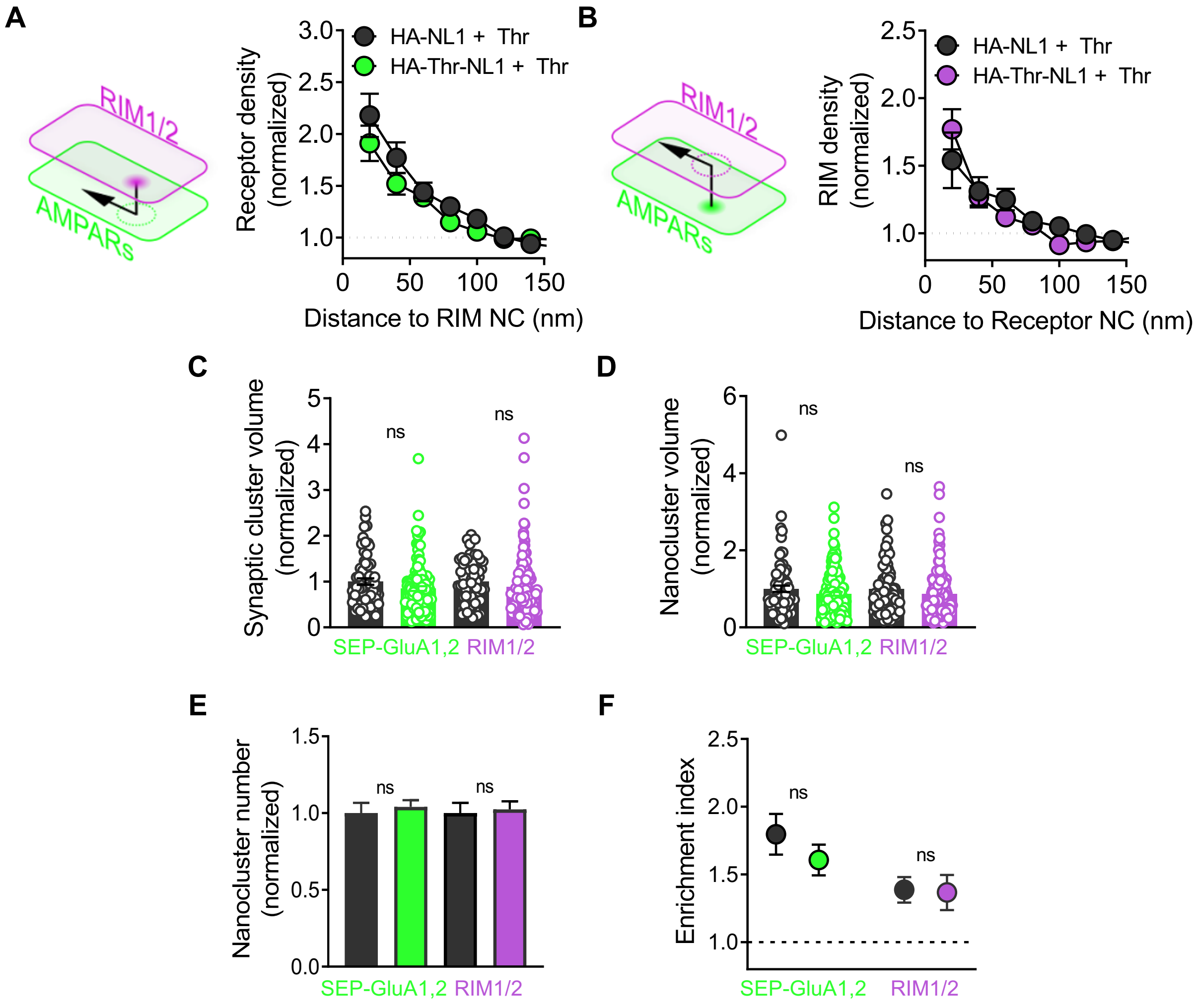


**Fig. S10.** Accompanies Fig 6. **(A)** Left, schematic demonstrating the measurement of AMPAR density across from RIM nanodomains in neurons transfected with SEP-GluA1,2 and HA-NL1-Thr or HA-NL both treated with thrombin. Right, quantification of receptor density as a function of distance to the RIM nanocluster. **(B)** Left, schematic demonstrating the measurement of RIM density across from AMPAR nanodomains in neurons transfected with SEP-GluA1,2 and HA-NL1-Thr or HA-NL both treated with thrombin. **(C)** Quantification of normalized synaptic cluster volume, **(D)** normalized nanocluster volume, **(E)** nanocluster number, and **(F)** enrichment indices. Data are presented as mean ± SEM, * $p \leq 0.05$, ** $p \leq 0.01$, *** $p \leq 0.001$. Mann Whitney rank-sum test was performed for C-F.
